# Fine-scale sampling uncovers the complexity of migrations in 5th-6th century Pannonia

**DOI:** 10.1101/2022.09.26.509582

**Authors:** Deven N. Vyas, István Koncz, Alessandra Modi, Balázs Gusztáv Mende, Yijie Tian, Paolo Francalacci, Martina Lari, Stefania Vai, Péter Straub, Zsolt Gallina, Tamás Szeniczey, Tamás Hajdu, Rita Radzevičiūtė, Zuzana Hofmanová, Sándor Évinger, Zsolt Bernert, Walter Pohl, David Caramelli, Tivadar Vida, Patrick J. Geary, Krishna R. Veeramah

## Abstract

As the collapse of the Western Roman Empire accelerated during the 4th and 5th centuries, arriving “barbarian” groups began to establish new communities in the border provinces of the declining (and eventually former) empire. This was a time of significant cultural and political change throughout not only these border regions but Europe as a whole.^1,2^ To better understand post-Roman community formation in one of these key frontier zones after the collapse of the Hunnic movement, we generated new paleogenomic data for a set of 38 burials from a time series of three 5th century cemeteries^3–5^ at Lake Balaton, Hungary. We utilized a comprehensive sampling approach to characterize these cemeteries along with data from 38 additional burials from a previously published mid-6th century site^6^ and analyzed them alongside data from over 550 penecontemporaneous individuals^7–19^. The range of genetic diversity in all four of these local burial communities is extensive and wider ranging than penecontemporaneous Europeans sequenced to date. Despite many commonalities in burial representation and demography, we find that there were substantial differences in genetic ancestry between the sites. We detect evidence of northern European gene flow into the Lake Balaton region. Additionally, we observe a statistically significant association between dress artefacts and genetic ancestry among 5th century genetically female burials. Our analysis shows that the formation of early Medieval communities was a multifarious process even at a local level, consisting of genetically heterogeneous groups.

## Results and Discussion

With the dissolution of the Western Roman Empire, the 5th century was a period of great political, cultural, and demographic change in Europe^1,2^. This was particularly true in the Middle Danube Region, which had long served as a frontier zone of the Roman Empire and also became a border zone after its division into the Western and Eastern Empires after 395 CE. With the abandonment of the Pannonian provinces by the Roman civil and military administration, most likely in 433 CE, it had already lost its former political and military importance. The region’s subsequent development was first determined by the period of Hunnic rule in the middle of the 5th century, after which it came under the influence of various “barbarian” groups (Goths, Heruls, Langobards, etc.). These changes resulted in the transformation of settlement structures and patterns, the appearance of new material culture, and the continuous emergence of new communities that founded small burial sites, quite unlike the large late Roman cemeteries of the 4th century, which could sometimes contain thousands of graves.^1,2^

We previously performed a comprehensive paleogenomic characterization of Szólád, a Langobard-period cemetery found on the southern shore of Lake Balaton, Hungary, dating to the middle of the 6th century.^6^ We demonstrated that the community was organized primarily around a large, three-generation male kindred enriched for genomic ancestry found in modern northern Europeans, and that it was possible to distinguish at least two groups of burials with different genomic ancestry that differed with regard to grave structure, the dress accessories, and other grave goods.

In this study we present the analysis of paleogenomic data derived from a dense spatial and temporal sampling approach^13,20,21^ for 38 individuals from three earlier cemeteries near Szólád that date to the middle and second half of the 5th century CE (Fonyód, Hács, and Balatonszemes). This allowed us to investigate whether the genetic heterogeneity observed at Szólád represented a long-standing post-Roman structure in the region or was the result of 6th-century population movements as described in the written sources. In particular we were interested in a) to what extent the continuous emergence of these short-lived communities was the result of extensive human mobility in the region and the appearance of new population groups, as attested by the written sources and b) what role biological relatedness played in the construction of social kinship and organization of these post-Roman communities.^6,22^ In other words, did genetic variation correspond to significant social and cultural differences in burial customs and spatial organization of the sites?

### 5th century Pannonia: Fonyód, Hács, & Balatonszemes

Fonyód, Hács, and Balatonszemes (Supp. Section S1) are located within 18 km of each other on elevated loess ridges close to the southern shore of Lake Balaton. They were chosen for analysis based on their geographical proximity to Szólád, and are considered fairly typical of the region and this period. They chronologically represent a time transect of the second part of the 5th century, with Fonyód dated to the middle third, Hács to the second half, and Balatonszemes to the end of the 5th century (**Fig. 1**). Their dating is based on certain jewelry types (brooches, pins, earrings, etc.) and tools (‘nomadic mirrors’ of the Čmi-Brigetio type, double-sided combs)^3–5,23,24^. All three sites represent small (19, 29, and 19 graves respectively), probably short-lived rural communities, with a substantial bias towards adult female burials (i.e., 20 adult females, 4 adult males, 17 subadults). Hács and Balatonszemes are also very similar in terms of burial representation, as artefacts are mostly found in female burials in contrast to male burials which are generally poorly furnished.^4,5,24^ Artificial cranial deformation (henceforth ACD) is also observable in all sites but it is only prevalent in Fonyód (found in 7/11 preserved crania), while the other two sites only contained single examples (Supp. Section S2). The sites also show differences in terms of spatial organization of the burials, with Fonyód being unique^24^ due to its six grave clusters lying at roughly equal distances of 50–60 m from each other (**Fig. 2**).

**Figure 1:**
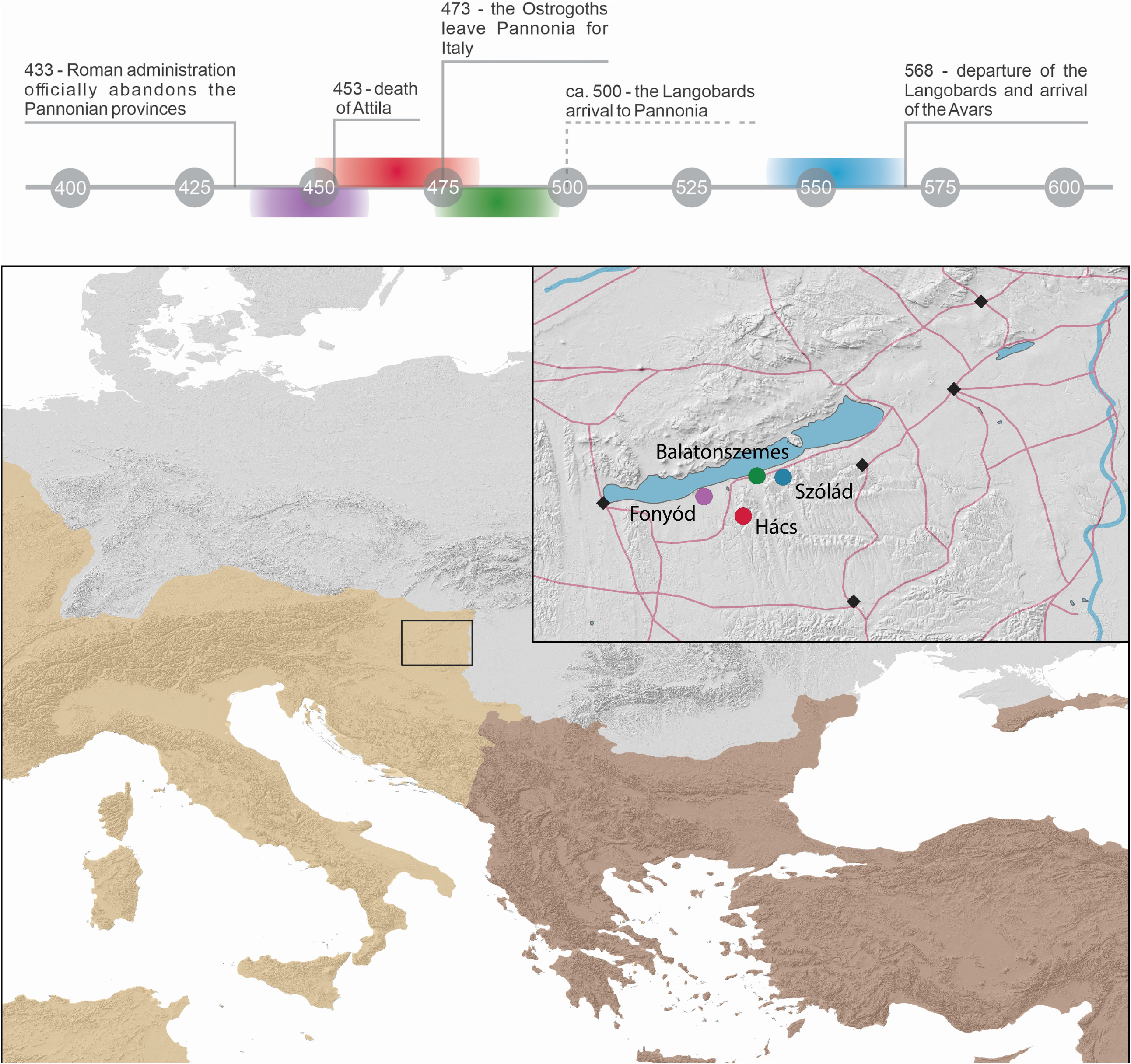
Location and chronological position of the four investigated sites within the (Western) Roman Empire. Roman roads are indicated with red lines, while late Roman inner fortresses of the Lake Balaton area are marked with black. Map created with QGiS v3.22.1 and resources used from Ancient World Mapping Center. “Coastline, River, Inland water, Open water, Roman road”. <http://awmc.unc.edu/wordpress/map-files/> [Accessed: January 07, 2022].^62^

**Figure 2:**
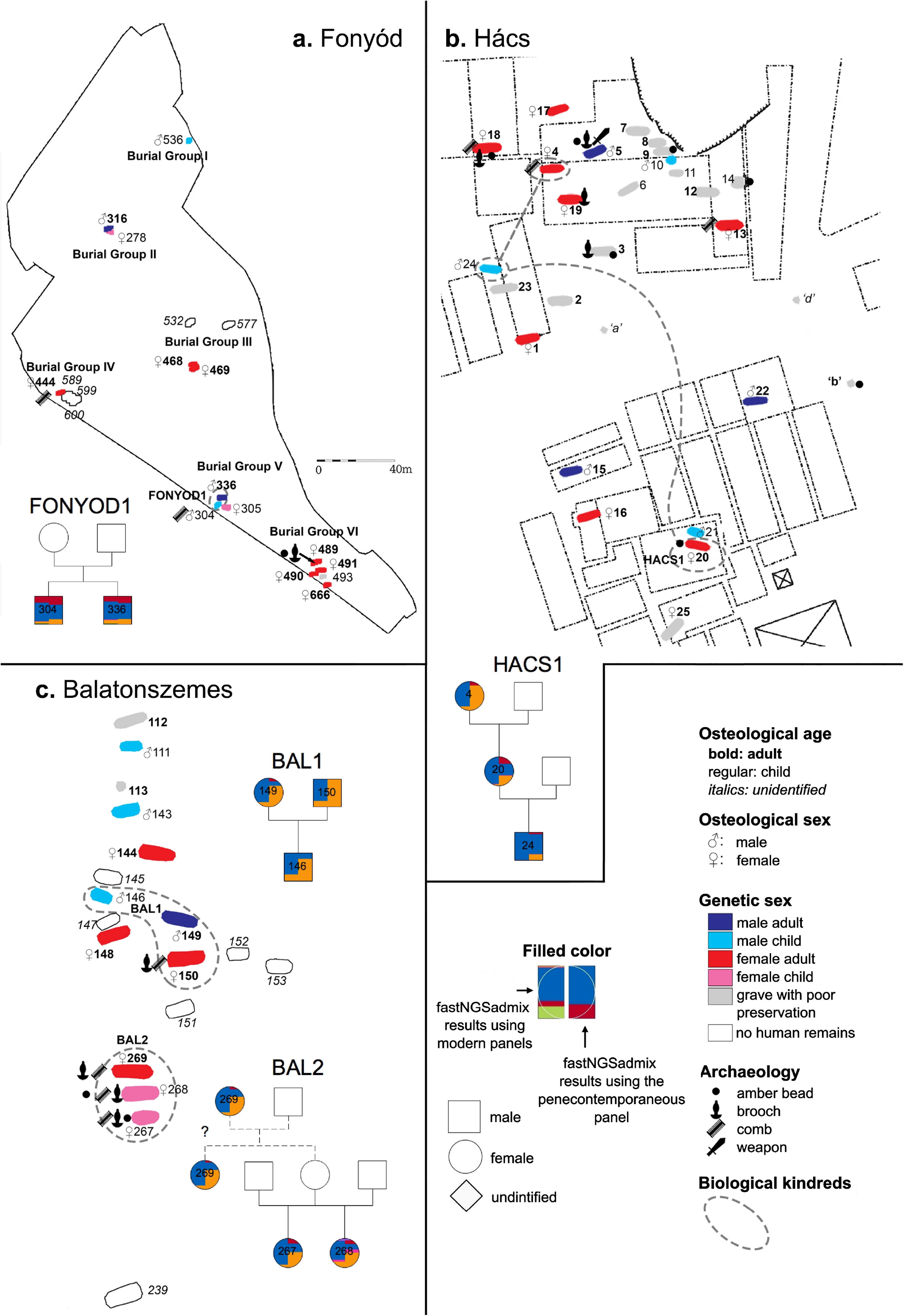
Cemetery maps of the three sites. Graves are identified based on age, (osteological and genetic) sex, and sampling status as well as the presence of amber beads, brooches, double-and single-sided combs, and weapons. Dotted outlines indicate biological relatedness. Pedigrees of the biologically related individuals are overlayed on the cemetery maps with colors corresponding to fastNGSadmix results from Figure 4. **a**. Cemetery map of Fonyód. Due to the dimensions of the cemetery, graves sizes are not to scale with the size of the cemetery. This figure was modified from Gallina and Straub.^24^ **b**. Cemetery map of Hács. This figure was modified from Kiss 1995.^5^ **c**. Cemetery map of Balatonszemes. Bal_26 is not included in this figure, as this individual was buried 200 meters west of the main site. This figure was modified from Miháczi-Pálfi 2018.^63^

### Genome sequencing

From Fonyód, we obtained genomic data from 13 of 14 graves with human remains, from Hács from all 14 accessible burials, and from Balatonszemes 11 of 13 graves with human remains. Altogether 38 samples were newly sequenced and analyzed (Supp. Section S3). Ten samples underwent whole genome sequencing (WGS) (coverage mean 8.36×, range 5-11×), and the remaining 28 underwent genome-wide SNP capture sequencing for ∼1.2 million SNPs (1240K)^25–27^ (coverage mean 1.47×, range 0.02-3.46×). Mitochondrial contamination rate medians^28^ were 1-3% for all individuals, while nuclear contamination point estimates^29^ from the 14 males were between 0.4% and 2% (Table S1). In addition, 14 new samples from Bardonecchia and Torino Lavazza (5th-8th century Italy) underwent 1240K capture with coverages from 0.39-3.85×, while one from Bardonecchia underwent WGS (Bard_T1, coverage 6.54×).

### Temporal Variation in Population genetic structure in the 5th century

A principal component analysis (PCA) of ancient individuals against modern reference individuals using a Procrustes-transformation technique demonstrated that all 5th century Lake Balaton individuals along with 6th-century Szólád possessed genomic ancestry that reflected primarily modern European genetic diversity (n=69 total) (**Fig 3a**, Supp Fig S1). In addition, we analyzed a penecontemporary (c. 4-8th century CE) set of 492 comparative individuals (477 previously published alongside 15 newly produced from Bardonecchia and Torino Lavazza) sampled from across Europe (see Table S2 and Supp. Section S4)^6–12,30^. Confidence ellipses of PC1 and PC2 coordinates demonstrate that while the penecontemporary regions cluster primarily with modern individuals from the same geographic regions (i.e., they are localized) (**Fig 3b**), our 5-6th century Lake Balaton individuals have a much more diverse genomic ancestry, encompassing a range reflecting the entire north to south axis of modern European genetic variation in the POPRES dataset^31^ (though not western or eastern Europe) (**Fig 3a**). While all Lake Balaton individuals were sampled from a single region of 200 km^2^ over a ∼100 years, the penecontemporary populations generally encompass much larger geographic regions and wider time frames (and thus we would expect more genetic diversity): this suggests that that this part of central Europe experienced particularly high rates of gene flow during the 5-6th centuries. We explicitly modeled gene flow in a spatial context across Europe using our Lake Balaton and penecontemporary individuals using FEEMS^32^ and indeed found a strong rate of gene flow between the Lake Balaton region and populations across Northern Europe ∼1,000km away (**Figs 3d, S2**)

**Figure 3:**
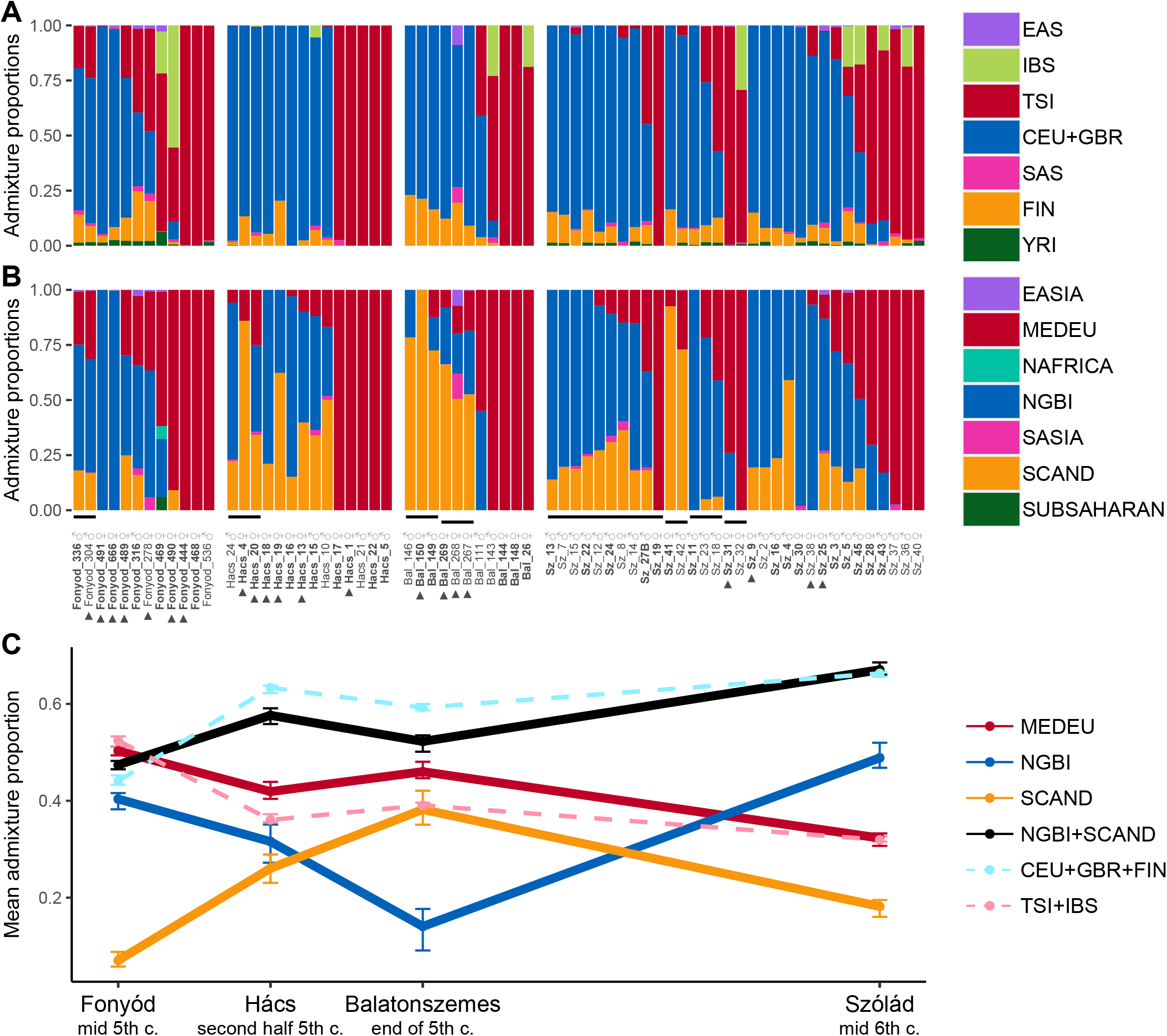
Procrustes PCA of 561 individuals^6–12,30^ transformed on to a PCA with 1,385 modern European individuals from the POPRES dataset^31^ using pseudohaploid genotype calls from 328,670 SNPs with individuals. Penecontemporaneous reference individuals are colored in pastel colors based on their region, while modern individuals from POPRES are colored in grey. **a**. PCA results from the Lake Balaton communities are plotted with covariance confidence ellipses with radii corresponding to 1.5 standard deviations overlaying the PCA. **b**. PCA results from the penecontemporary populations are plotted with similar ellipses. **c**. PCA zoomed in on the northern cluster of the PCA with polygons identifying the lack of overlap between the 5th century Lake Balaton communities. **d**. a FEEMS plot estimating gene flow between the 4th-8th century communities analyzed in the PCA with all four Lake Balaton communities analyzed together. Bluer colors indicate more migration, while brown indicate less/no migration.

To further explore the structure within our Lake Balaton cemeteries, we also used a model-based clustering method (fastNGSadmix)^33^ to characterize the genomic ancestry of each individual using 1000 Genomes Project (1000G)^34^ populations as possible sources. In addition, given the localized geographic structure observed in the PCA for the penecontemporary Europeans, we constructed a panel of penecontemporary reference individuals with a similar geographic distribution to the 1000G populations (see Supp. Section S5)^7,9–12,14–19^. The two reference sets yielded qualitatively similar patterns (**Figs 4a-b and S3**), with almost all individuals possessing some combination of primarily Tuscan (TSI), Central European and Great Britain (CEU+GBR), and Finnish (FIN) modern ancestry or Mediterranean (MEDEU), northern Germany and British (NGBI), and Scandinavian/Estonian (SCAND) penecontemporary ancestry. However, all three 5th-century cemeteries have distinct profiles with regard to the relative proportions of these components, with the penecontemporary analysis showing these differences most clearly. Most notably there is a clear progressive significant increase in SCAND ancestry and corresponding decrease in NGBI ancestry with time for the 5th century sites (Fonyód, Hács, Balatonszemes SCAND 0.07>0.26>0.38, NGBI 0.40<0.31<0.14, **Fig 4c**).

**Figure 4:**
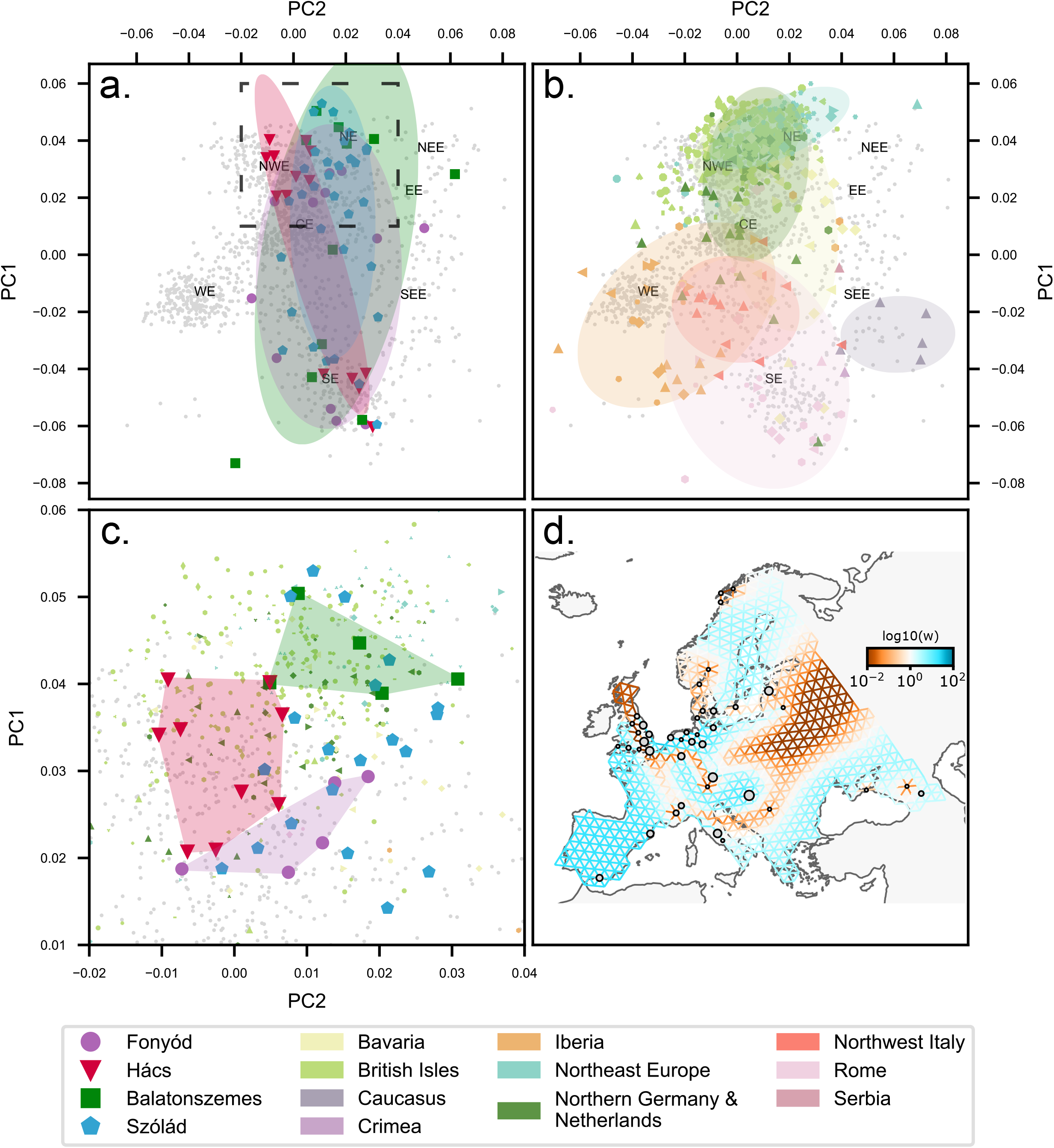
Supervised ancestry proportions from individuals from Fonyód, Hács, Balatonszemes, and Szólád. The ♂ and ♀ symbols identify genetic males and genetic females, respectively, while triangles identify individuals buried with dress accessories (in the case of Fonyód, Hács, and Balatonszemes) or brooches specifically (in the case of Szólád). Proportions were estimated using fastNGSadmix in using **a**. using 1000 Genomes populations as references^34^ with CEU+GBR referring to a merger of “Northern Europeans from Utah” (CEU) and “British in England and Scotland” (GBR), FIN refers to “Finnish in Finland”, IBS refers to “Iberian populations in Spain”, and TSI refers to “Tuscans from Italy”; EAS refers to the East Asian super-population, SAS refers to the South Asian super-population, and YRI refers to “Yoruba in Ibadan, Nigeria”. **b**. penecontemporaneous individuals to form reference populations^7,9–12,14–19^ with MEDEU referring to individuals from Italy and Iberia (Mediterranean Europe), NGBI refers to individuals from what are now northern Germany and Britain, SCAND refers to individuals from what are now Scandinavia/Estonia, EASIA refers to individuals from what is now Hanben, Taiwan, NAFRICA refers to individuals from what is now Sudan, SASIA refers to individuals from Roopkund Lake in what is now India, and SUBSAHARAN refers to individuals from sub-Saharan Africa. Individuals with <0.1× coverage were excluded from this analysis. Individuals are sorted based on increasing MEDEU and then by decreasing NGBI. **c**. presents a line graph showing the change in ancestry proportions through time over the four sites.

Fonyód, which dates to the peak of the Hunnic power in the region, differs markedly from the later two 5th century sites from both an archaeological and genomic perspective. Its unique spatial structure suggests that it is a burial site of a short-lived coexistence of Hunnic period groups, part of them practicing ACD, a tradition that is considered a foreign phenomenon in Pannonia and less prevalent in the later sites.^35^ Genomically, Fonyód also possesses significantly (as assessed by non-overlapping 95% CIs) more Mediterranean ancestry than the other Lake Balaton sites and is somewhat southern shifted on the PCA. Overall, the site shows more non-European diversity, with two individuals possessing ∼6% Asian ancestry and one individual 12% African ancestry. Thus the genomic profile of Fonyód may reflect heterogeneity of the late Roman period and/or recent influx from the East.

Both Hács and Balatonszemes are enriched for individuals with very high proportions of northern European ancestry (NGBI+SCAND) that are not as prominent at Fonyód (**Fig 3c**), potentially indicating the arrival of new groups to the region consistent with written records describing the emergence of new ‘barbarian’ powers after the fall of the Hunnic empire, in the second half of the 5th century. The high level of gene flow between Lake Balaton individuals and northern Europe is only seen in our FEEMS analysis when examining these later two communities, while the use of the earlier Fonyód site results in a barrier to gene flow to this region and greater gene flow with southern Europe (**Fig S2**). However because of variation in the relative SCAND/NGBI components, individuals with higher PC1 values (i.e,. more northern European ancestry) show no overlap in the PCA between the two sites (**Fig 3c**), with Hács angling towards the modern northwestern Europe and Balatonszemes northeastern Europe. These differences could indicate that groups from similar, yet distinguishable, sources from northern Europe arrived into the region in multiple waves during the second half of the 5th century. On the other hand, the presence of individuals with high amounts of MEDEU ancestry is a constant in all four sites and they show overlap in the PCA, suggesting this may represent a more stable local genomic signature during this entire period.

Interestingly, Szólád demonstrates a profile that encompasses genomic variation observed in all three 5th-century sites, albeit it has a much larger sample size. Northern European ancestry is even more prominent than at Hács and Balatonszemes, driven primarily (but not totally) by a large nine member pedigree identified previously. The individuals in this pedigree along with most other northern-like individuals lack the high SCAND component observed in Balatonszemes (qualitatively resembling those from Fonyód and Hács instead, the three notable exceptions being two second degree relatives, Sz_41 and Sz_42, with 92% and 73% SCAND ancestry respectively and Sz_4 with 59%), suggesting no major direct continuity between these two sites. Szólád is notable for strontium isotope data suggesting most adults at the site were non-local, regardless of genomic ancestry. However our data make clear that the major patterns of genetic ancestry observed at Szólád were already established during the 2nd half of the 5th century in and around the Lake Balaton area. Thus, this community could have formed from the existing diverse regional pool of genetic variation established ∼50 years earlier, rather than just being the result of the arrival of a new population group, i.e., the Langobards, as interpreted by both historical and archaeological research.^36–39^

### Biological relatedness and social relationships

The small burial groups at Fonyód were previously interpreted as signs of strong social ties^24^ and Hács and Balatonszemes have been described as family cemeteries.^5,23,40^ We used lcMLkin to identify close biological relatives in all three 5th-century sites. At all three sites, biological kindreds (i.e., sets of closely biologically related individuals) consisted of only a few very close, first- and second-degree relatives, mostly on the mother’s side (**Fig 2**, Table S3). This is in stark contrast to Szólád where the cemetery was organized largely around male biological relatives with a large extended pedigree. Biologically related individuals at Fonyód and Balatonszemes were buried in close proximity to each other and in the latter case their connection is also clearly reflected by the similarities in the burial customs suggesting that these connections held meaningful social values. (Supp. Section S6). However, the relatively low number of biologically related individuals, the small size of the kindreds and the lack of biological relations in most burial groups—probably also influenced by the short occupation period of the sites and the low number of buried individuals—suggest that other factors may have had a major influence on the formation of these 5th century communities.

To examine to what extent the broader biological backgrounds of individuals (as determined by genomic ancestry) beyond close biological kinship were acknowledged as meaningful social ties, we also compared burial customs demonstrating variation with the POPRES PCA PC1 and PC2 coordinates using a logistic regression framework (Table S4). We observed a strong significant association between the presence of various jewelry types and dress accessories (i.e., polyhedral earrings, various brooch types, bracelets, amber beads, etc.) characteristic of the 5^th^-century female burials^41,42^ and genetic variation (p=0.008, n=20 for adults, p=0.002, n=23 for adults and children). This appears to be primarily driven by most individuals with dress accessories possessing high amounts of northern European ancestry (13/16 are majority SCAND/NGBI) (**Fig 4, S4**), and as such, site-specific regressions are only significant for Hács (p=0.01, n=8) and Balatonszemes (p=0.008, n=7) but not Fonyód (p=0.08, n=8) (despite all three tests having similar power; Table S4). This results points to individuals with northern European genetic backgrounds being treated differently in death by their respective communities, perhaps marking cultural and/or social differences between locals and more recent migrants in the region. Though close biological relatives were rare, it is also noteworthy that all biological kindreds involved individuals with predominantly northern genomic ancestry, maybe reflecting another aspect of this biologically-structured social organization.

While Szólád shows a similar genomic profile to the three 5th-century sites, it is strikingly different in terms of funerary practices, spatial organization and demographics. There was no significant association between genetic variation and brooches—an artefact that in various forms and types is present in the 5th-century sites as well as the 6th-century Szólád—from genetically female burials (including and excluding subadults) (p<0.56, n=11), suggesting that social and economic differences were no longer formulated along fault lines of genetic background in this community. Whether this weakening of the association between this artefact and genomic ancestry reflects a more general shift in social kinship practices (at least with respect to women) since the 5th century and the establishment of Langobard rule is difficult to conclude with just our data, as even amongst other 6th century Lake Balaton sites Szólád is considered somewhat unique, with the vast majority of adult and adolescent males (15/19) being buried with weapons.^36,43^ The importance of a northern genomic background appears to still be manifested in this community socially, but rather through the dominance of the cemetery by a spatially clustered and primarily male kindred.

## Conclusion

Early medieval history has played a major role in the formation of modern Europe. Ethnonationalist rhetoric in the 19th and 20th centuries allowed a link to be made between historical peoples from various written sources and modern nation states.^44,45^ The key to this linkage was the idea of a common ethnic and cultural heritage from these presumed ancestral and homogenous groups after the fall of the Roman Empire, though recent historical, archaeological, and anthropological studies have found this to be a vast oversimplification, with the material culture demonstrating significant complexity across the early Middle Ages. Strikingly, rather than homogeneity, our three post-Roman 5th-century sites from Lake Balaton exhibit considerable genomic diversity compared to penecontemporaneous Europe. This region experienced particular high levels of gene flow during this period, and there are significant shifts in genomic ancestry through our 5th century time transect over a period of only ∼50 years that indicate migration into the region from various sources, probably from areas in northern Europe, consistent with the continuously changing political landscape of the period described in the historical record.^46,47^ As these post-Roman communities formed from mixtures of locals and migrants, social ties between individuals of similar genomic backgrounds appear to have still been maintained to some extent, but these links could shift rapidly rather than being enduring states of social organization, and the importance of close biological relatedness varied considerably. Given the immense complexity observed in just these four small cemeteries, it is clear that comprehensive fine-grained spatiotemporal genomic sampling will be critical to unpack the processes that underlie the subsequent development of modern Europe.

## STAR Methods

### Processing of novel genomic dataset

Petrous bone samples were collected from individuals from Fonyód, Hács, and Balatonszemes (as well as two penecontemporaneous Italian reference sites, Bardonecchia and Torino Lavazza), except Fonyod_304 and Fonyod_305 where tibial fragments were collected instead. The bone samples were powdered in a clean room exclusively dedicated to ancient DNA analysis, and a silica-based protocol was used for DNA extraction and purification^48^. Illumina libraries were created and processed for all DNA samples. For Hács, Balatonszemes, Bardonecchia, and Torino Lavazza exploratory shotgun sequencing was conducted, and libraries underwent a target enrichment for ∼1.2 million genome-wide SNPs (1240K) before sequencing, or a direct whole deep genome sequencing (WGS) according to endogenous DNA preservation; alternatively, for Fonyód, all libraries directly went to 1240K target enrichment, omitting shotgun sequencing (see Supp. Section S3).

BAM files from these novel sites were called using in-house scripts (www.github.com/kveeramah). Non-UDG treated data was processed using a caller that incorporates damage patterns from mapDamage, while UDG treated data was instead processed using an indent caller that disregarded the first and last five bases of any given read. Output VCF files include diploid genotypes as well as genotype likelihoods. For detailed information about these sites, see Supp. Section S1. For detailed information about DNA extraction and processing, see Supp. Section S3. Information about all individuals newly processed and analyzed for this paper are presented in Table S1.

### Mitochondrial and Y chromosome analyses

For mitochondrial analysis, BAM files from the four Balaton region sites were filtered down to reads aligning to the mitochondrial genome as fastq files using samtools. Mitochondrial genomes from individuals with >1500 mitochondrial reads were assembled using Mapping Iterative Assembler ^49^ (which was designed for ancient mitochondrial genome assembly). Haplotype assignment was made using MitoTool 1.1.2.^50^ For further mtDNA analysis, see Supp. Section S7.

For the Y chromosome, we restricted the analysis to 1240K Y chromosome SNPs among 1240K-captured males but analyzed the whole chromosome for the four WGS males. Phylogenetic position of each Y chromosome SNP was ascertained based on published databases (https://isogg.org/)^34,51–53^ and used to identify NRY haplogroups.

### Comparative dataset

We analyzed our newly sequenced data alongside comparative datasets.^6–12,14–19,30^ All individuals analyzed had date ranges overlapping with the 4th-8th centuries CE with the exception of four individuals from Olalde et al.^11^, where date cut-offs were broadened to increase the number of Iberian reference individuals. For all comparative individuals, coverage of the autosomal 1240K SNPs was calculated using gatk DepthOfCoverage,^54^ and (with the exception of four individuals from Szólád) individuals with <0.1× coverage were excluded from our dataset and all analyses. In most cases, we used the BAMs published in the study, but in some cases we used BAMs that were re-processed in the Veeramah lab from published fastq files using the same type of pipeline as our novel data. BAMs were called using in-house genotype callers consistent with our novel data. Information on all comparative individuals analyzed are presented in Table S2.

All individuals (both reference and novel) were called for genotypes from the whole genome, imputed for all diploid, dinucleotide sites within the 1000 Genomes Project Phase 3 v5a VCF files using GLIMPSE v1.1^55^ (using the 1000 Genomes Project data as the imputation reference)^34^, and filtered down to the 1240K positions.

### Biological relatedness assessment

We used the software package lcMLkin to identify close biological relatedness between the individuals from all four Balaton sites (as well as Bardonecchia and Torino Lavazza).^56^ All individuals were included regardless of level of coverage. We used a modified version of lcMLkin that included external allele frequency data, similar to Amorim et al.^6^ Analyses were performed using genotype likelihood data from 1,079,996 autosomal 1240K sites as input. We used allele frequency data from the 1000 Genomes CEU and TSI populations (as well as the merger of the two, CEU_TSI)^34^. We ignored all relationships between low-coverage individuals with minimal common SNP coverage (i.e., less than 5,000 shared SNPs in an lcMLkin analysis) as these are likely spurious (not real) relationships.

### Principal component analysis

We analyzed the data from the Balaton region sites alongside data from 492 penecontemporaneous comparative individuals. PCAs were conducted to place ancient individuals one-by-one onto a background of modern individuals. A Procrustes transformation was used to merge 561 ancient Europeans onto a single PCA (with a background of modern individuals). We used modern European populations from the POPRES dataset^31^ as imputed by Veeramah et al.^30^ We conducted analyses using all overlapping 328,670 SNPs (Supp. Section S4). For each Balaton region site as well as for each geographic region, we also calculated covariance confidence ellipses based on Pearson correlation coefficients; ellipses were given radii corresponding to 1.5 standard deviations.

Logistic regressions testing association of PCA results with dress accessories and ACD were calculated using R v4.1.2^57^ and plotted using the ggplot2^58^ and cowplot packages; power analyses were conducted using the WebPower package.^59^ Hosmer and Lemeshow’s R^2^ and chi-square p values for each regression were calculated within R.

### Gene flow modeling analysis

We used FEEMS^32^ in order to model gene flow in a spatial context across Europe using our Lake Balaton and penecontemporary individuals. We used GLIMPSE imputations described above. FEEMS has a native imputation feature for missing data, but this simply relies on the mean allele frequencies across individuals for a given SNP, and does not account for low coverage. FEEMS also relies on population allele frequencies, which are likely to be estimated more accurately for low coverage data than individual genotypes. Of the 566 4-8^th^ century Europeans available to us, 26 were removed due to inferred 1-3^rd^ degree relatedness between other individuals due to the reliance on unbiased population frequencies. We further set a maximum of n=100 for individuals represented from a particular macro-region (for example British Isles) in order to mitigate as much as is possible against biases that result from oversampling in one region and undersampling in others. For macro-regions with more than 100 individuals, we randomly chose 100 representatives. We filtered out SNPs with a minor allele frequency less than 0.1 in the CEU 1000 Genomes populations^34^ and performed linkage disequilibrium filtering using plink using --indep-pairwise 50 5 0.2.^60^ Thus the final dataset consisted of 371 4-8^th^ century individuals from across Europe with imputed diploid genotypes at 76,669 SNPs. We note that this compares favorably to the example use case of FEEMS in Marcus et al.^32^ using 111 gray wolves from across North America genotyped at 17,729 SNPs (though that dataset has a more even geographic spread and relies on true diploid genotypes rather than imputation). An approximate polygon of Europe encompassing the geographic location of our sampleset was obtained using https://www.birdtheme.org/useful/v3tool.html. FEEMS was run under default parameters under three different lambda values (20, 2, 0,2). Decreasing lambda decreases the smoothing but can lead to overfitting, but we found a lambda of 0.2 to provide the best resolution of meaningful geographic barriers, with larger numbers leading to overly smooth surfaces that were not informative), and thus we provide visualizations from using this value. We conducted this analysis using all Lake Balaton individuals simultaneously (Fig 3c) and for each of the four populations individually (Fig S3).

### Model-based clustering analyses

We conducted supervised ancestry clustering analyses using fastNGSadmix.^33^ We constructed seven reference panels of imputed penecontemporaneous individuals representing Mediterranean Europe^7,9–12,14–19^ (Italy, Iberia; MEDEU, n=40), Northern Germany/Britain (NGBI, n=40), Scandinavia/Estonia (SCAND, n=40), East Asia (EASIA, n=16), South Asia (SASIA, n=17), North Africa (NAFRICA, n=20), and sub-Saharan Africa (SUBSAHARAN, n=8). We also conducted analyses using modern 1000 Genomes Project populations, CEU+GBR, FIN, IBS, TSI, YRI, EAS, and SAS; as per Amorim et al.^6^, we merged the CEU and GBR populations into a single population, as the two populations are not properly distinguishable by these types of analyses.

We generated the Beagle PL files for the genotype likelihoods used by fastNGSadmix for all individuals with coverage ≥0.1× for 1,091,054 autosomal 1240K sites using vcftools.^61^ FastNGSadmix analyses were run for each individual one-by-one. Each individual analysis was run 50 times with 10 bootstraps per run for penecontemporaneous individuals and 100 bootstraps for Balaton individuals; the run with the greatest (i.e., least negative) likelihood was used for analysis. We also conducted a version using 1000 Genomes Project data^34^ to supervise the analyses (see Supp. Section S5).

## Data availability

Our newly generated sequence data from 53 individuals are available from the NCBI Sequence Read Archive (SRA) database under accession PRJNA811958 for 1240K data from Fonyod, Hács, and Balatonszemes; under accession PRJNA812074 for WGS data from Hács and Balatonszemes; and under PRJNA936918 for 1240K and WGS data from Torino Lavazza and Bardonecchia. We accessed the POPRES (Population Reference Sample) dataset collected and published by Nelson et al. ^31^ from dbGaP (accession phs000145.v4.p2).

## Acknowledgements

This project has received funding from the European Research Council (ERC) under the European Union’s Horizon 2020 research and innovation programme (grant agreement n° 856453 ERC-2019-SyG), from the Anneliese Maier Research Award of the Alexander von Humboldt Foundation, from the Max Planck Society, from the German Federal Ministry for Education and Research, from the Swedish Riksbankens Jubieleumfond, by the Czech Grant Agency (GACR 21-17092X), from the Gerard B. Lambert Foundation, from the Institute for Advanced Study Director’s Office, and from the Italian Ministry of Education, University and Research (project “Dipartimenti di Eccellenza 2018–2022” and PRIN2017 grants no. 20177PJ9XF). We thank Szilvia Honti, Péter Németh, and the Rippl-Rónai Museum, Kaposvár for providing access to the material from Balatonszemes; the Hungarian Natural History Museum, Budapest for providing access to the anthropological material from Fonyód and Hács; and Caterina Giostra, Luisella Pejrani Baricco, and the late Elena Bedini for access to the material from Bardonecchia and Torino Lavazza. We also thank Johannes Krause for funding and performing all SNP capture and sequencing.

## Supplemental Table Titles

**Supplemental Table S1**. Information about newly sequenced data

**Supplemental Table S2**. Information of comparative ancient dataset

**Supplemental Table S3. l**cMLkin results using three different modern allele frequency panels for pairs sharing at least 5000 SNPs (sorted by CEU+TSI Pi-hat value).

**Supplemental Table S4**. Results from Logistic Regression analyses

## References

1. Hardt, M. (2011). Pannonien im Spannungsfeld zwischen Römer- und Völkerwanderungszeit – eine geschichtliche Einführung. In Keszthely-Fenékpuszta im Kontext spätantiker Kontinuitätsforschung zwischen Noricum und Moesia Castellum Pannonicum Pelsonense 2., O. Heinrich-Tamáska, ed. (Leidorf), pp. 15–28.

2. Vida, T. (2011). Die Zeit zwischen dem 4. und dem 6. Jahrhundert im mittleren Donauraum aus archäologischer Sicht. In Römische Legionslager in den Rhein- und Donauprovinzen Abhandlungen., M. Konrad, ed. (Bayerische Akademie der Wissenschaften, phil.-hist. Klasse / NF 138), pp. 615–650.

3. Straub, P. (2008). Adalékok a Balaton környéki 5. századi temetők Felső-Duna vidéki kapcsolatához. Zalai Múzeum 17, 189–207.

4. Miháczi-Pálfi, A. (2017). Form-und herstellungstechnische Analyse der Bügelfibeln von Balatonszemes aus dem dritten Viertel des 5. Jahrhunderts. ANTÆUS: Communicationes Ex Instituto Archaeologico Academiae Scientiarum Hungaricae 35–36, 67–89.

5. Kiss, A. (1995). Das germanische Gräberfeld von Hács-Béndekpuszta (Westungarn) aus dem 5.--6. Jahrhundert. Acta Antiquae Academiae Scientiarum Hungaricae 36, 275–342.

6. Amorim, C.E.G., Vai, S., Posth, C., Modi, A., Koncz, I., Hakenbeck, S., La Rocca, M.C., Mende, B., Bobo, D., Pohl, W., et al. (2018). Understanding 6th-century barbarian social organization and migration through paleogenomics. Nat. Commun. 9, 3547.

7. Antonio, M.L., Gao, Z., Moots, H.M., Lucci, M., Candilio, F., Sawyer, S., Oberreiter, V., Calderon, D., Devitofranceschi, K., Aikens, R.C., et al. (2019). Ancient Rome: A genetic crossroads of Europe and the Mediterranean. Science 366, 708–714.

8. Damgaard, P. de B., Marchi, N., Rasmussen, S., Peyrot, M., Renaud, G., Korneliussen, T., Moreno-Mayar, J.V., Pedersen, M.W., Goldberg, A., Usmanova, E., et al. (2018). 137 ancient human genomes from across the Eurasian steppes. Nature 557, 369–374.

9. Gretzinger, J., Sayer, D., Justeau, P., Altena, E., Pala, M., Dulias, K., Edwards, C.J., Jodoin, S., Lacher, L., Sabin, S., et al. (2022). The Anglo-Saxon migration and the formation of the early English gene pool. Nature 610, 112–119.

10. Margaryan, A., Lawson, D.J., Sikora, M., Racimo, F., Rasmussen, S., Moltke, I., Cassidy, L.M., Jørsboe, E., Ingason, A., Pedersen, M.W., et al. (2020). Population genomics of the Viking world. Nature 585, 390–396.

11. Olalde, I., Mallick, S., Patterson, N., Rohland, N., Villalba-Mouco, V., Silva, M., Dulias, K., Edwards, C.J., Gandini, F., Pala, M., et al. (2019). The genomic history of the Iberian Peninsula over the past 8000 years. Science 363, 1230–1234.

12. Schiffels, S., Haak, W., Paajanen, P., Llamas, B., Popescu, E., Loe, L., Clarke, R., Lyons, A., Mortimer, R., Sayer, D., et al. (2016). Iron Age and Anglo-Saxon genomes from East England reveal British migration history. Nat. Commun. 7, 1–9.

13. Veeramah, K.R. (2018). The importance of fine-scale studies for integrating paleogenomics and archaeology. Curr. Opin. Genet. Dev. 53, 83–89.

14. Harney, É., Nayak, A., Patterson, N., Joglekar, P., Mushrif-Tripathy, V., Mallick, S., Rohland, N., Sedig, J., Adamski, N., Bernardos, R., et al. (2019). Ancient DNA from the skeletons of Roopkund Lake reveals Mediterranean migrants in India. Nat. Commun. 10, 1–10.

15. Prendergast, M.E., Lipson, M., Sawchuk, E.A., Olalde, I., Ogola, C.A., Rohland, N., Sirak, K.A., Adamski, N., Bernardos, R., Broomandkhoshbacht, N., et al. (2019). Ancient DNA reveals a multistep spread of the first herders into sub-Saharan Africa. Science 365, eaaw6275.

16. Sirak, K.A., Fernandes, D.M., Lipson, M., Mallick, S., Mah, M., Olalde, I., Ringbauer, H., Rohland, N., Hadden, C.S., Harney, É., et al. (2021). Social stratification without genetic differentiation at the site of Kulubnarti in Christian Period Nubia. Nat. Commun. 12, 1–14.

17. Skoglund, P., Thompson, J.C., Prendergast, M.E., Mittnik, A., Sirak, K., Hajdinjak, M., Salie, T., Rohland, N., Mallick, S., Peltzer, A., et al. (2017). Reconstructing Prehistoric African Population Structure. Cell 171, 59–71.e21.

18. Wang, K., Goldstein, S., Bleasdale, M., Clist, B., Bostoen, K., Bakwa-Lufu, P., Buck, L.T., Crowther, A., Dème, A., McIntosh, R.J., et al. (2020). Ancient genomes reveal complex patterns of population movement, interaction, and replacement in sub-Saharan Africa. Science Advances 6, eaaz0183.

19. Wang, C.-C., Yeh, H.-Y., Popov, A.N., Zhang, H.-Q., Matsumura, H., Sirak, K., Cheronet, O., Kovalev, A., Rohland, N., Kim, A.M., et al. (2021). Genomic insights into the formation of human populations in East Asia. Nature 591, 413–419.

20. Pohl, W., Krause, J., Vida, T., and Geary, P. (2021). Integrating Genetic, Archaeological, and Historical Perspectives on Eastern Central Europe, 400–900 AD: Brief Description of the ERC Synergy Grant – HistoGenes 856453. Historical Studies on Central Europe 1, 213–338.

21. Geary, P., and Veeramah, K. (2016). Mapping European population movement through genomic research. Mediev. Worlds medieval worlds, 65–78.

22. O’Sullivan, N., Posth, C., Coia, V., Schuenemann, V.J., Price, T.D., Wahl, J., Pinhasi, R., Zink, A., Krause, J., and Maixner, F. (2018). Ancient genome-wide analyses infer kinship structure in an Early Medieval Alemannic graveyard. Sci Adv 4, eaao1262.

23. Bondár, M., Honti, S., Márkus, G., and Németh, P.G. (2007). Balatonszemes-Szemesi berek. In Gördülő idő: régészeti feltárások az M7-es autópálya Somogy megyei szakaszán Zamárdi és Ordacsehi között, K. Belényesy, S. Honti, and V. Kiss, eds. (Somogy Megyei Múzeumok Igazgatósága), pp. 123–135.

24. Gallina, Z., and Straub, P. (2014). Fonyód–Vasúti-dűlő 2 (Mérnöki-telep) kora népvándorlás kori sírjai. In AVAROK PUSZTÁI Régészeti tanulmányok Lőrinczy Gábor 60. születésnapjára, A. Alexandra, B. Csilla, and T. Attila, eds. (MTA BTK MOT), pp. 213–233.

25. Fu, Q., Meyer, M., Gao, X., Stenzel, U., Burbano, H.A., Kelso, J., and Päbo, S. (2013). DNA analysis of an early modern human from Tianyuan Cave, China. Proc Natl Acad Sci USA 110, 2223–2227.

26. Haak, W., Lazaridis, I., Patterson, N., Rohland, N., Mallick, S., Llamas, B., Brandt, G., Nordenfelt, S., Harney, E., Stewardson, K., et al. (2015). Massive migration from the steppe was a source for Indo-European languages in Europe. Nature 522, 207–211.

27. Mathieson, I., Lazaridis, I., Rohland, N., Mallick, S., Patterson, N., Roodenberg, S.A., Harney, E., Stewardson, K., Fernandes, D., Novak, M., et al. (2015). Genome-wide patterns of selection in 230 ancient Eurasians. Nature 528, 499–503.

28. Renaud, G., Slon, V., Duggan, A.T., and Kelso, J. (2015). Schmutzi: estimation of contamination and endogenous mitochondrial consensus calling for ancient DNA. Genome Biol. 16, 224.

29. Korneliussen, T.S., Albrechtsen, A., and Nielsen, R. (2014). ANGSD: Analysis of Next Generation Sequencing Data. BMC Bioinformatics 15, 356.

30. Veeramah, K.R., Rott, A., Groß, M., van Dorp, L., López, S., Kirsanow, K., Sell, C., Blöcher, J., Wegmann, D., Link, V., et al. (2018). Population genomic analysis of elongated skulls reveals extensive female-biased immigration in Early Medieval Bavaria. Proc Natl Acad Sci USA 115, 3494–3499.

31. Nelson, M.R., Bryc, K., King, K.S., Indap, A., Boyko, A.R., Novembre, J., Briley, L.P., Maruyama, Y., Waterworth, D.M., Waeber, G., et al. (2008). The Population Reference Sample, POPRES: a resource for population, disease, and pharmacological genetics research. Am. J. Hum. Genet. 83, 347–358.

32. Marcus, J., Ha, W., Barber, R.F., and Novembre, J. (2021). Fast and flexible estimation of effective migration surfaces. Elife 10, e61927.

33. Jørsboe, E., Hanghøj, K., and Albrechtsen, A. (2017). fastNGSadmix: admixture proportions and principal component analysis of a single NGS sample. Bioinformatics 33, 3148–3150.

34. 1000 Genomes Project Consortium, Auton, A., Brooks, L.D., Durbin, R.M., Garrison, E.P., Kang, H.M., Korbel, J.O., Marchini, J.L., McCarthy, S., McVean, G.A., et al. (2015). A global reference for human genetic variation. Nature 526, 68–74.

35. Hakenbeck, S. (2018). Infant Head Shaping in Eurasia in the First Millennium AD. In The Oxford Handbook of the Archaeology of Childhood, S. Crawford, D. M. Hadley, and G. Shepherd, eds. (Oxford University Press), pp. 483–504.

36. Alt, K.W., Knipper, C., Peters, D., Müller, W., Maurer, A.-F., Kollig, I., Nicklisch, N., Müller, C., Karimnia, S., Brandt, G., et al. (2014). Lombards on the Move – An Integrative Study of the Migration Period Cemetery at Szólád, Hungary. PLoS One 9, e110793.

37. Jarnut, J. (1982). Geschichte der Langobarden (W. Kohlhammer).

38. Pohl, W., and Erhart, P. (2005). Die Langobarden : Herrschaft und Identität (Verlag der Österreichischen Akademie der Wissenschaften).

39. von Freeden, U., Vida, T., and Skriba, P. (2007). Ausgrabung des langobardenzeitlichen Gräberfeldes von Szólád, Komitat Somogy, Ungarn: Vorbericht und Überblick über langobardenzeitliche Besiedlung am Plattensee. Germania 85, 359–384.

40. Hakenbeck, S.E., Evans, J., Chapman, H., and Fóthi, E. (2017). Practising pastoralism in an agricultural environment: An isotopic analysis of the impact of the Hunnic incursions on Pannonian populations. PLoS One 12, e0173079.

41. Rácz, Z. (2016). Zwischen Hunnenund Gepidenzeit. Frauengräber aus dem 5. Jahrhundert im Karpatenbecken. Acta Archaeologica Academiae Scientiarum Hungaricae 67, 301–359.

42. Rácz, Z. (2020). Who Were the Gepids and Ostrogoths on the Middle Danube in the 5th Century? An Archaeological Perspective. In The Migration Period between the Oder and the Vistula (2 vols) (Brill), pp. 771–789.

43. Barbiera, I. (2005). Changing lands in changing memories : migration and identity during the Lombard invasions (All’Insegna del giglio).

44. Geary, P.J. (2002). The Myth of Nations: the Medieval Origins of Europe (Princeton University Press).

45. Fehr, H. (2010). Germanen und Romanen im Merowingerreich: Frühgeschichtliche Archäologie zwischen Wissenschaft und Zeitgeschehen: Ergänzungsbände zum Reallexikon der Germanischen Altertumskunde (Gruyter, Walter de, & Co.).

46. Heather, P.J. (2005). The fall of the Roman Empire : a new history of Rome and the Barbarians (Macmillan).

47. Meier, M. (2021). Geschichte der Völkerwanderung : Europa, Asien und Afrika vom 3. bis zum 8. Jahrhundert n.Chr (C.H.Beck).

48. Dabney, J., Knapp, M., Glocke, I., Gansauge, M.-T., Weihmann, A., Nickel, B., Valdiosera, C., García, N., Päbo, S., Arsuaga, J.-L., et al. (2013). Complete mitochondrial genome sequence of a Middle Pleistocene cave bear reconstructed from ultrashort DNA fragments. Proc Natl Acad Sci USA 110, 15758–15763.

49. Briggs, A.W., Good, J.M., Green, R.E., Krause, J., Maricic, T., Stenzel, U., Lalueza-Fox, C., Rudan, P., Brajković, D., Kucan, Ž., et al. (2009). Targeted Retrieval and Analysis of Five Neandertal mtDNA Genomes. Science 325, 318–321.

50. Fan, L., and Yao, Y.-G. (2013). An update to MitoTool: using a new scoring system for faster mtDNA haplogroup determination. Mitochondrion 13, 360–363.

51. Francalacci, P., Morelli, L., Angius, A., Berutti, R., Reinier, F., Atzeni, R., Pilu, R., Busonero, F., Maschio, A., Zara, I., et al. (2013). Low-pass DNA sequencing of 1200 Sardinians reconstructs European Y-chromosome phylogeny. Science 341, 565–569.

52. Karmin, M., Saag, L., Vicente, M., Wilson Sayres, M.A., Järve, M., Talas, U.G., Rootsi, S., Ilumäe, A.-M., Mägi, R., Mitt, M., et al. (2015). A recent bottleneck of Y chromosome diversity coincides with a global change in culture. Genome Res. 25, 459–466.

53. Poznik, G.D., Henn, B.M., Yee, M.-C., Sliwerska, E., Euskirchen, G.M., Lin, A.A., Snyder, M., Quintana-Murci, L., Kidd, J.M., Underhill, P.A., et al. (2013). Sequencing Y chromosomes resolves discrepancy in time to common ancestor of males versus females. Science 341, 562–565.

54. McKenna, A., Hanna, M., Banks, E., Sivachenko, A., Cibulskis, K., Kernytsky, A., Garimella, K., Altshuler, D., Gabriel, S., Daly, M., et al. (2010). The Genome Analysis Toolkit: a MapReduce framework for analyzing next-generation DNA sequencing data. Genome Res. 20, 1297–1303.

55. Rubinacci, S., Ribeiro, D.M., Hofmeister, R.J., and Delaneau, O. (2021). Efficient phasing and imputation of low-coverage sequencing data using large reference panels. Nat. Genet. 53, 120–126.

56. Lipatov, M., Sanjeev, K., Patro, R., and Veeramah, K.R. (2015). Maximum Likelihood Estimation of Biological Relatedness from Low Coverage Sequencing Data. bioRxiv, 023374. 10.1101/023374.

57. R Core Team (2022). R: A language and environment for statistical computing R Foundation for Statistical Computing. Vienna, Austria.

58. Wickham, H. (2009). ggplot2: Elegant Graphics for Data Analysis (Springer, New York, NY).

59. Zhang, Z., and Yuan, K.-H. (2018). Practical statistical power analysis using Webpower and R (Isdsa Press).

60. Purcell, S., Neale, B., Todd-Brown, K., Thomas, L., Ferreira, M.A.R., Bender, D., Maller, J., Sklar, P., de Bakker, P.I.W., Daly, M.J., et al. (2007). PLINK: a tool set for whole-genome association and population-based linkage analyses. Am. J. Hum. Genet. 81, 559–575.

61. Danecek, P., Auton, A., Abecasis, G., Albers, C.A., Banks, E., DePristo, M.A., Handsaker, R.E., Lunter, G., Marth, G.T., Sherry, S.T., et al. (2011). The variant call format and VCFtools. Bioinformatics 27, 2156–2158.

62. QGIS Development Team (2022). QGIS Geographic Information System (QGIS Association).

63. Miháczi-Pálfi, A. (2018). A balatonszemesi 5. századi temető kisleletei. Anyagközlés és elemzés [Kleinfunde des Gräberfeldes von Balatonszemesaus dem 5. Jahrhundert. Katalog und Bewertung der Funde]. In "Hadak útján.” A népvándorláskor fiatal kutatóinak XXVI. konferenciája. Gazdaság – kereskedelem – kézművesség (26th Conference of Young Scholars on the Migration Period. Economy – Trade – Craftsmanship) Dissertationes Archaeologicae Supplementum., Z. Rácz, I. Koncz, and B. Gulyás, eds. (Institute of Archaeological Sciences, Eötvös Loránd University, Budapest), pp. 129–161.

