## Supplemental for "Fine-scale sampling uncovers the complexity of migrations in 5th-6th century Pannonia"

**Supplemental Text**

### S1 Description of Archaeological sites

#### S1.1 Fonyód, Hács, & Balatonszemes

All three sites are located on elevated loess ridges close to the southern shore of Lake Balaton, an area that became increasingly important for the Roman military and civil administration with the founding of a series of inner fortresses in the 3rd-4th centuries. These inner fortresses remained in use at least until the middle of the 5th century, sometimes even after the Roman abandonment of the area.[1–3] These sites represent small, rural communities with similarities in both age and sex distributions and burial customs. However, they show differences in terms of spatial organization of the burials and due to chronological differences in archaeological material (Fig. 2). The three sites chronologically cover the second part of the 5th century with Fonyód dated to the middle third, Hács to the second half, and Balatonszemes to the end of the 5th century.

The most interesting feature of the Fonyód site is its unique structure: 19 graves form six clusters lying at roughly equal distances of 50–60 m from each other (Fig. 2a). Some of the burial groups lay along the boundaries of the investigated area, so the site cannot be considered completely excavated. The 19 graves contained 14 individuals with 7 adult females, 2 adult males, 5 children, and 5 graves without any biological remains. Out of the 11 well-preserved skulls at least 7 showed signs of artificial cranial deformation (henceforth ACD), that includes both adult male, female, and child burials. For the full discussion of the ACD found at Fonyód, see Section S2. Curiously enough, adult male and adult female burials never occurred in the same cluster; however, given the low number of burials, no far-reaching conclusions can be drawn from this pattern. Most of the graves are simple pits with two exceptions found in Group VI: the pit of Fonyod_489 followed the contours of the human body, while Fonyod_666 was a side niche grave. The site is dated to the middle of the 5th century, to the peak of the Hunnic movement based on the presence of certain artefact types, such as the so called 'nomadic mirrors' of the Čmi-Brigetio type from Fonyod_444 and Fonyod_489 or the silver pins from Fonyod_491.

The cemetery at Hács contains at least 29 burials with a very unbalanced sex ratio (13 identifiable adult females and only 3 adult males) (Fig. 2b). The burials form several clusters, but since the cemetery has not been completely excavated and at least four graves were destroyed, its exact layout cannot be reconstructed. Burials were more concentrated in the northern section, while they were more dispersed in the southern part, lying at distances of 5–10 m from each other or forming pairs. The site is dated to the second half of the 5th century based on the Béndekpuszta-type brooches found in Hacs_19 and Hacs_20, while polyhedral earrings and double-sided combs give a wider chronological frame.[4] The most remarkable finds of the cemetery are the delicate lead sheet fragments—possibly used as an amulet—bearing a text inscribed with the Gothic uncial and Gothic cursive script, probably found in the heavily disturbed Grave 5.[5]

At Balatonszemes, 19 graves were unearthed with five adult females, one adult male, two unidentifiable adults, five children, and six pits lacking any human remains (Fig. 2c). A large grave cluster with twelve graves is arranged into a fairly irregular north to south row. Three burials (Bal_267, Bal_268, Bal_269) lying some 7-8 m to its south form a separate cluster. Yet another burial (Bal_26) was uncovered some 200 m west of the other graves, whose association with the cemetery remains unclear, even though the grave goods indicate contemporaneity.[6] While most burials barely contained any artefacts, four burials (two adult females and two girls including one girl with ACD) were remarkably richly furnished. Archaeological dating is again based on female jewelry and brooches found in the four richly furnished burials are the best chronological indicators pointing to the end of the 5th century.[7,8]

#### S1.2 Torino Lavazza

To the north of ancient Turin, on the other side of the Dora river in the close vicinity of a long-known, late Roman necropolis used between the 1st to 4th centuries, an early Christian funerary complex including burials, mausolea, and a church with single nave and semi-circular apse have been discovered in 2013.[9] The complex developed as an organic extension of the late Roman necropolis with the first burials and several mausoleums appearing at the site during the 3rd-4th centuries. This phase is dated based on late Roman artefacts (most notably glass) of the period, but also based on the structure and masonry technique of the buildings. One of the mausolea—an apsidal hall—was probably turned into a funerary church sometimes later attested by the presumed later dating of burials both in the apse and the hall.

Later, in the second half of the 4$^{th}$ century or during the 5$^{th}$ century a large building with a single hall and semi-circular apse facing to the west, was erected above the foundations of the earlier mausolea in the northern part of the site. The building has been interpreted as a suburban funerary basilica. Funerary use of the site continued well into the 5$^{th}$ century with burials appearing inside and around the church suggesting an intense and systematic funerary use. Most graves of this phase are similar to the late Roman tombs: rectangular pits with masonry walls (re)using Roman bricks, while their bottom is often paved with joined bricks or tiles. Due to the lack of artifacts in the burials and the absence of epigraphic evidence, burials of this phase are dated based on their stratigraphy, their relative position to the earlier and later burials. The sampled burials belong to this phase, however due to the lack of absolute dating methods, their dating can only be given as 5$^{th}$-7$^{th}$ centuries. Later phases also show the systematic and accurate exhumation of the skeletal remains of the masonry tombs that suggest the partial abandonment of the site and degradation of the buildings probably used as stone quarries, although there is some evidence in written records that suggests the survival of the church into the 11$^{th}$-12$^{th}$ centuries.

**S1.3 Bardonecchia**

In 2005 a small cemetery was excavated near the today's commune of Bardonecchia, 90 kms west from Turin, Italy at an intersection of multiple valleys and surrounded by mountains.[10,11] Altogether 16 graves—four of them without any human remains, while others containing multiple individuals—came to light, but as it is located on a steep, south-oriented slope, it is probable that an unknown number of graves were destroyed by erosion. The east-west oriented graves are simple pits with occasional stone slab or stone lining around the edges, and they form irregular north-south rows

From male burials belt buckles and mounts, knives, a double-sided comb, and a scramasax came to light. Female artefact types, such as jewelry, are only known from the burial of a child, possibly a young girl, while other female burials lacked any grave goods or dress accessories. Based on the artifacts the site can be dated from the second half of the 6th and to the 7th century, but the site might have survived into the 8th century, as most of the burials are undatable due to the lack of grave goods. C14 dating of multiple burials attest the wide dating range between the 6th and 8th centuries.

Based on the abundance of males and much fewer females, the presence of a weapon and the relatively large number of fractures and cranial trauma, it has been suggested that the community using the Bardonecchia cemetery might have had a military role in controlling the region and the mountain passes leading through the Susa valley.

#### S2 Artificial cranial deformation

Artificial cranial deformation (ACD) was found at all three of the 5th century sites we studied, but it was most prominent at Fonyód. Out of the 11 well-preserved skulls at Fonyód, at least 7 showed signs of ACD, including adult male, female, and child burials. Based on the techniques (i.e. placement and number of bandages) used, three different types of artificially deformed skulls were observed: Fonyod_304, 305, 336, 490, 491 constitute Type 1; Type 2 is Fonyod_489 and Type 3 is Fonyod 666 (Fig. S2).

The first type is *moderate fronto-occipital ACD with two bandages*, where the first pressure bandage had been tied around the forehead and the occipital bone and was adjusted more tightly than the second one. The second bandages were situated behind the bregmatic region and either under the mandible or around the nuchal region. This is the most common type at the site (Fonyod_304, 305, 336, 490, 491). Fonyod_468 shows similar characteristics, but the deformation is so mild that it can be the result of natural variation as well.

The second type is *severe oblique cranial deformation with one bandage*: the modification was made by applying one bandage fastened between the frontal and the occipital region producing an obliquely conical skull shape. This type is only present in Fonyod_489.

The third type is *severe oblique artificial cranial deformation with two bandages*. The modification is the result of a strongly fastened bandage encircling the frontal and occipital bone resulting in an oblique elongation and conical shape of the skull. The second bandage had been placed in a vertical position around the bregmatic region and either under the mandible or around the nuchal region. This type is only present in Fonyod_666.

While it was possible to distinguish three types of ACD based on the techniques used, the second (Fonyod_489) and third (Fonyod_666) types resulted in very similar, elongated skulls that would have been indistinguishable to people living at the time, so only two distinctive forms (three including not-deformed) can be recognized. Additionally, Fonyod_468 has marks that might indicate a slight modification, but any such deformation would have been unrecognizable during their lifetime.

There is a clear connection between the spatial organization of the site and ACD. All individuals with recognizable deformed skulls were found in Burial group V and VI, where all preserved skulls showed signs of this tradition. Fonyod_489 and Fonyod_666 are both found in Burial group VI. Individuals from Burial group I, II, III, or IV did not show any signs of ACD with the exception of the female with the possibly slightly modified skull from Fonyod_468.

We conducted a logistic regression comparing ACD to our PCA results among the 10 sequenced individuals with well-preserved skulls and coverage greater than 0.1×; we coded Fonyod_468 as without ACD. Across all comparisons (POPRES-Full, POPRES-Tv, AADR WE), we found significant associations between PCA results and ACD

At the other two fifth century sites ACD is less prominent, but is still present in Hács (Hacs_23) and Balatonszemes (Bal_268). Hacs_23 is an adult female (who was not sequenced), and Bal_268 is a young girl. Both of these individuals had deformed skulls with similar shapes to type 2 and/or 3. Another interesting factor is that in our fastNGSadmix analyses, Bal_268 (unlike all other individuals from Balatonszemes and Hács) has significant proportions of Asian ancestry components in both the fastNGSadmix analyses using 1000 Genomes Project and penecontemporaneous reference panels (Fig. 5). Interestingly, her maternal half-sister (Bal_267) lacks both ACD and these modern/historic Asian components.

#### S3 Ancient DNA Lab Work and Sequence Processing

##### S3.1 Overview

Molecular work for the specimens from Hács and Balatonszemes and from Fonyód was carried out at the University of Florence and the Institute of Archaeogenomics, Eötvös Loránd Research Network in Budapest (respectively). Initial steps were identical between all three sites.

Work was conducted in dedicated ancient DNA clean room facilities, applying strict criteria to prevent contamination during all experimental procedures.[12,13] Moreover, blank controls were processed along with the samples during DNA extractions and libraries preparation to monitor for contamination in reagents. Before sampling the bone powder, the outer layer of the temporal and petrous bones was brushed with disposable tools and irradiated by ultraviolet light (254 nm) for 30 minutes, to remove external contaminants. To maximize the recovery of well-preserved endogenous DNA, powder was collected from the densest part of the pars petrosa (inner ear) as described in Pinhasi et al.[14] (with the exceptions of Fonyod_304 and Fonyod_305 where tibial samples were collected instead), using a low-speed micro drill equipped with a disposable disk saw and dental burs. Subsequent steps differ between data from Hács, Balatonszemes, Torino Lavazza, and Bardonecchia (which were processed in 2018) and data from Fonyód (which were processed in 2021).

For Hács, Balatonszemes, Torino Lavazza, and Bardonecchia, all 1240K library capture was conducted at the Max Planck Institute for the Science of Human History (MPI-SHH) in Jena, Germany, while libraries were prepared at the University of Florence. For Fonyód, all libraries and 1240K captures were prepared at the Max Planck Institute for Evolutionary Anthropology (MPI-EVA) Leipzig, Germany.

##### S3.2 Hács, Balatonszemes, Torino Lavazza, and Bardonecchia

Libraries from Hács, Balatonszemes, Torino Lavazza, and Bardonecchia were sequenced in 2018, while the libraries from Fonyód were sequenced separately in 2021. For the sites processed in 2018, all sequencing was performed at the New York Genome Center (NYGC). Initially, 40 double-stranded libraries were shotgun sequenced at NYGC using an Illumina benchtop sequencer (MiSeq). All 40 libraries passed screening and were chosen for further sequencing; the libraries that had the most reads mapping to the human reference genome (GRCh37) were chosen for whole genome sequencing (WGS), whereas the other libraries underwent 1240K capture sequencing for 1.24 million SNPs. In total, ten were chosen for WGS, nine from Hács and one from Balatonszemes. For 1240K sequencing from Hács (n=5) and Balatonszemes (n=10), double-stranded libraries were sequenced using paired-end 125bp sequencing on an Illumina HiSeq 2500 sequencer at NYGC. For WGS sequencing, double-stranded libraries were sequenced using single-end 100bp sequencing on an Illumina NovaSeq S4 sequencer at NYGC with each WGS library sequenced over four different flow cell lanes.

Approximately 50 mg of bone powder per sample were used for DNA extraction at the Laboratory of Molecular Anthropology and Paleogenetics at the University of Florence. Extraction was conducted using a silica-based protocol that allows ancient DNA molecules to be efficiently recovered even if highly fragmented.[15] For these four sites, Illumina sequencing libraries without enzymatic damage repair were prepared from 20 uL of each extract using a double-strand and double-indexing protocol optimized for ancient samples.[16,17] After quality control on Agilent 2100 Bioanalyzer (DNA 1000 chip), libraries were pooled in equimolar amounts and sent to the New York Genome Center (NYGC) for sequencing where all 40 double-stranded libraries were shotgun sequenced using an Illumina benchtop sequencer (MiSeq) along with libraries from other sites (not discussed here). All libraries passed screening.

Genomic libraries were subsequently sent to the Max Planck Institute for the Science of Human History (MPI-SHH) where libraries were enriched for endogenous human DNA using capture probes targeting ~1.24 million SNPs (1240K)[18–20] using a protocol similar to Fonyód described below. These 1240K libraries were sent to NYGC for paired-end 125bp sequencing on an Illumina HiSeq 2500 sequencer. Nine libraries from Hács, one from Balatonszemes, and one from Bardonecchia (Hacs_1, Hacs_4, Hacs_5, Hacs_10, Hacs_13, Hacs_17, Hacs_18, Hacs_21, Hacs_22, Bal_268, and Bard_T1), which performed best in the screening, also had WGS libraries prepared at the University of Florence. WGS libraries were sent to NYGC for single-end 100bp sequencing on an Illumina NovaSeq S4 sequencer with each WGS library sequenced over four different flow cell lanes.

**S3.3 Fonyód**

For Fonyód, all available genomic libraries underwent 1240K capture sequencing without any screening phase. These libraries were single-stranded[21,22] and were UDG treated in a manner with results functionally similar to partial UDG treatment or UDG-half.[23] These libraries were sequenced using single-end 76bp sequencing on an Illumina HiSeq 4000 sequencer at the MPI-EVA.

DNA extraction and subsequent steps of sample preparation were performed in the Ancient DNA Core Unit of the Max Planck Institute for Evolutionary Anthropology (MPI-EVA), Leipzig, Germany. DNA was extracted from between 21.8 to 35.7 mg of sample material using the same silica-based method optimized for the recovery of short DNA fragments.[15] Briefly, lysates were prepared by adding 1 ml of extraction buffer (0.45 M EDTA, pH 8.0, 0.25 mg/ml proteinase K, 0.05% Tween-20) to the sample material in 2.0-ml Eppendorf Lo-Bind tubes and rotating the tubes at 37°C for approximately 16 hours.[15,24] Using an automated liquid handling system (Bravo NGS Workstation B, Agilent Technologies), DNA was purified from 150 µl lysate using silica-coated magnetic beads and binding buffer D as described in Rohland et al.[24] Elution volume was 30 µl. Extraction blanks without sample material were carried alongside the samples during DNA extraction.

DNA libraries were prepared from 30 µl extract using an automated version of single-stranded DNA library preparation[25] described in detail in Gansauge et al.[26] *E. coli* Uracil-DNA-glycosylase and *E. coli* endonuclease VIII were added during library preparation to remove uracils in the interior of molecules. Libraries were prepared from both the sample DNA extracts and the extraction blanks, and additional negative controls (library blanks) were added. Library yields and efficiency of library preparation were determined using two quantitative PCR assays.[26] The libraries were amplified and tagged with pairs of sample-specific indices using AccuPrime Pfx DNA polymerase as described in Gansauge et al.[26] Amplified libraries were purified using SPRI technology [27] as described in Gansauge et al.[26]

Sample and control libraries were enriched for endogenous human DNA using capture probes targeting ~1.24 million SNPs (1240K).[18–20] Two consecutive rounds of in-solution hybridization capture were performed using the method of Fu et al. [18] automated on the Bravo NGS workstation B. Pools of up to 24 libraries (also of different projects) were created and sequenced in single-end mode for 76 cycles on 3 lanes of 2 independent runs of an Illumina HiSeq 4000 sequencer.

**S3.4 Genome assembly, Genotype calling, and data quality**

Sequencing data was processed based on a pipeline from Kircher [28]. Reads were trimmed and merged (when necessary), mapped to GRCh37 using samtools [29], duplicate reads were marked using Picard Tools [30], and reads less than 30bp long were filtered out. All untrimmed reads were excluded from processing and analysis, as these reads are more prone to contain contaminant sequences. As WGS libraries were sequenced over four lanes, data from each lane were initially processed separately but BAM files were merged prior to marking duplicate reads. We used mapDamage 0.3.3 to assess postmortem DNA damage patterns [31].

We calculated coverage for all individuals for the autosomal 1240K SNPs (as well as for the mitochondrial genome) using gatk v4.2 DepthOfCoverage; for novel WGS data, we also calculated full autosomal coverage using gatk v3.3 DepthOfCoverage in multi-threaded operation.[30] We calculated the number of autosomal 1240K SNPs covered by at least one read with mapping and base quality scores greater than or equal to 30 in each BAM file using samtools depth.[29] Individuals' genetic sex was identified using the Sex.DetERRmine pipeline to determine relative X and Y chromosome coverage (https://github.com/TCLamnidis/Sex.DetERRmine). Subsequently, we used Angsd to estimate nuclear contamination rates of genetic males based on the hemizygous X chromosome.[32] We also used Schmutzi to estimate mitochondrial contamination rates for all individuals.[33] For seven individuals, Schmutzi failed to complete (Bard_T2, Bard_T6, To_Lav_T10US67_ind2, To_Lav_T1US6, To_Lav_T2US16, To_Lav_T37US60, To_Lav_T38US344), likely due to insufficient non-duplicate reads, similar to the results from Gnecchi-Ruscone et al.[34] However, six of these seven were males with successful Angsd results suggesting low contamination (thus only To_Lav_T38US344 lacks any contamination estimates) (Table S1).

#### S4 Pseudohaploid PCA using POPRES dataset

We conducted a principal component analysis comparing the individuals from our four sites to modern populations. For this analysis, we converted our diploid 1240K VCF files to pseudohaploid transposed Plink datasets using a custom, in-house script. Heterozygous genotype calls were converted to calls for the allele with greater allele depth; when both alleles had the same depth, an allele was chosen at random.

For our comparative datasets, we used modern European populations from the POPRES dataset [35] in its imputed form from Veeramah et al.[36] All analyzed POPRES genotypes were also made pseudohaploid by randomly choosing one allele for all heterozygous genotypes.

The autosomal 1240K sites and imputed POPRES dataset had an overlap of 328,670 SNPs. We used an automated script to use smartPCA[37,38] to perform PCAs on the POPRES dataset including one ancient individual at a time (https://github.com/ShyamieG/). We then used an in-house script to conduct a Procrustes transformation, merging all ancient individuals (with 1240K coverage of at least 0.1×) onto a single principal component analysis (Figs. 3, S1, and S4). In our PCA plots for these analyses, we divide POPRES populations into regions (i.e., CE, EE, NE, NEE, NWE, SE, SEE, and WE) as demarcated in Veeramah et al.[36]

We also use normalized PC1 and PC2 results from the PCA for our logistic regressions for testing association between artefacts (or ACD) and genetic ancestry (Table S4).

#### S5 Model-based clustering analyses using modern and penecontemporary references

In our study, we took a new approach to supervised model-based clustering analyses. Instead of solely using modern data for reference panels, we also conducted analyses using penecontemporary reference populations. All penecontemporary individuals used in reference panels were imputed for the entire genome (based on the 1000 Genomes Project Phase 3 v5a vcf files)[39] using Glimpse v1.1[40] and then filtered down to the autosomal 1240K positions. Due to a bias in coverage in favor of the more northern reference populations in our set (Table S2), imputation was necessary. Preliminary, non-imputed analyses found considerable bias towards the northern components as there was substantially more genotype data from the higher coverage data from northern Europe.[41–43] Furthermore, this bias also made it impossible to filter imputed genotypes based on genotype probabilities (as these are closely linked to coverage).

For panel construction, we attempted to construct penecontemporaneous reference panels[41–51] to mirror the 1000 Genomes Project populations (as used by Amorim et al.[52]); however, we could not create a working historic analog for the IBS population. Thus, we merged the penecontemporaneous Italian and Iberian groups into a single MEDEU panel. For the three European panels (MEDEU, NGBI, and SCAND), we picked individuals with the highest coverage while preventing any kin pairs from being put into a panel together.[41–45] These three panels consisted of 40 individuals each. We also included two Asian and two African panels: EASIA (n=16 from what is now Hanben, Taiwan), NAFRICA (n=20 from the island of Kulubnarti in what is now Sudan), SASIA (n=17 from Roopkund Lake in what is now India), and SUBSAHARAN (n=8 from various sites).[46–51] The reduction in SUBSAHARAN size owes directly to the rarity of ancient DNA from sub-Saharan Africa, especially from our time window. Given the closer relationships between the European reference panels, we used larger sample sizes (n=40) for these panels, while using fewer for the non-European panels.

For each individual, we ran the fastNGSadmix analysis 50 times and selected the run with greatest (i.e., least negative) likelihood. For the selected run, we re-ran the analysis using the exact same random seed to replicate the final result and calculate 100 bootstraps for Lake Balaton individuals and ten bootstrap for penecontemporary individuals. This approach was necessary as calculating bootstraps 50 times per individual would have been too computationally taxing, and even more so on lower coverage individuals who take longer to converge. In Figure 4 of the main text, we have presented our analyses for the four Lake Balaton sites using both modern and penecontemporary panels. In Figure S4, we have also presented our results from the penecontemporaneous individuals. We found that for the most part the bootstrap analyses were largely consistent with the converged results. When we analyzed the reference individuals, we found essentially perfect recall (i.e., each individual is assigned the ancestry that they are a reference for) (Fig. S4b-c). It is important to note that while the reference panel was imputed, the data put into fastNGSadmix were raw, un-imputed genotype likelihoods; thus, this supports the accuracy of our imputations.

We also conducted a version of the analysis where we used separate IBERIA (n=25) and ITALY (n=40) reference panels (Fig. S4c). The IBERIA panel was smaller due to fewer available individuals; outside of Olalde et al.[45], we were unable to find additional published, penecontemporary Iberians and even extended our 4th-8th century date cut-offs to include I8339, I10866, I10892, and I10895 (Table S2). Based on analyzing bootstraps the IBERIA component frequently appeared and disappeared within individuals between bootstraps. We particularly found these cases in Fonyod_304, Fonyod_316, and Fonyod_489 as well as several individuals from the datasets from Veeramah et al.[36] and Gretzinger et al.[42]. We also noticed the lack of IBERIA in individuals such as Bard_T11, FN2, I3056, and IND006, that were clustered with Iberians in our PCAs and contained significant IBS proportions in the 1000 Genomes Project analysis (Figs. 3 and S4). Based on these findings, we found that the IBERIA component could not be fully differentiated, and thus opted to merge IBERIA and ITALY into MEDEU for our primary analyses.

#### S6 Description of biological kindreds found at Fonyód, Hács, and Balatonszemes

A major aim of this study was to examine to what extent genetic connections between individuals observed at these post-Roman 5th century Pannonian sites were known and acknowledged by members of their communities and thus could be understood as a component of social kinship. Biological relatedness might form the basis of or play an important role in the construction of social kinship, but social kinship can also be organized based on shared space, imagined or real descent, economical connection, social agreement, etc. as it is an outcome of social actions, developed and expressed through culturally defined social practices, including funerals.[53–55] While there is a slight chronological difference between the three sites) the sites show similar characteristics in demography, funerary customs, and burial representation, and at all three sites, biological kindreds consisted of very close, first- and second-degree relatives, mostly on the mother's side. This is in stark contrast to Szólád where the cemetery was organized largely around male biological relatives with a large extended pedigree.

In Fonyód and Balatonszemes biologically related individuals are buried near each other suggesting that biological relatedness likely had social value to the community. Except for the three related females buried in richly furnished burials (Bal_267, Bal_268, and Bal_269) in Balatonszemes, these connections are not observable in the grave goods, probably as a result of differences in burial representation between the different genders. Adult male and child burials contain very few if any artefacts (notable exceptions to the latter are Bal_267 and Bal_268), while social differences are more clearly expressed in female burials.[56,57] At Hács, in contrast, the lack of spatial clustering and similarities in terms of burial representation may initially suggest a reduced role for biological relatedness among the three individuals directly related along the maternal line, a connection that is especially hard to hide among members of a small community. Previous stable isotope results[58] suggest that Hacs_4 was most likely raised locally, but probably gave birth to her daughter elsewhere, as the $^{87}Sr/^{86}Sr$ ratio of Hacs_20 falls completely outside of the regional values of the Lake Balaton area. Ancestry clustering analysis shows that across the generations of this kindred that SCAND proportions are being replaced with NGBI proportions (**Figs. 2b and 4**), suggesting that. Hacs_4 and Hacs_20 had children with men who had predominantly NGBI ancestry. This signal of replacement is not visible when using the 1000 Genomes Project reference panel.The fact that both women came back to this community later during their lives and were buried at Hács indicates that their connection to the community never ceased to exist and that their biological relatedness nevertheless reflected social ties as well.

#### S7 Analyses of Mitochondrial DNA Variation

Outside of haplogroup identification, we restricted our mtDNA analyses to the HVS-I region, as our mitogenome assemblies are low coverage and contain many ambiguous bases. We aligned all assembled mitogenomes and the rCRS sequencing using MAFFT v7.149b[59,60] using the G-INS-i option and extracted the bases corresponding to the HVS-I region based on the PhyloTree annotation[61] using Geneious Prime v. 2022.0.1 (https://www.geneious.com/). Close biological relatives were removed from the dataset (retaining only the highest coverage individual from each biological kindred, or removing Bal_146 in the case of BAL2), leaving behind 45 individuals. We conducted an AMOVA[62–64] and a population differentiation test[65,66] as well as calculated gene diversity[67] using the Arlequin 3.5 using the GUI application on Windows 10.[68]

Our AMOVA of the mitochondrial HVS-I indicates the mitochondrial variation is distributed within sites (not between)[62–64], and a test of population differentiation between all four Balaton region sites does not find any significant differences between the four sites[65,66]. Estimates of gene diversity of the HVS-I are different between the four sites (0.233 at Fonyód, 0.236 at Hács, 0.302 at Balatonszemes, and 0.145 at Szólád).[67]

#### Supplementary Figures

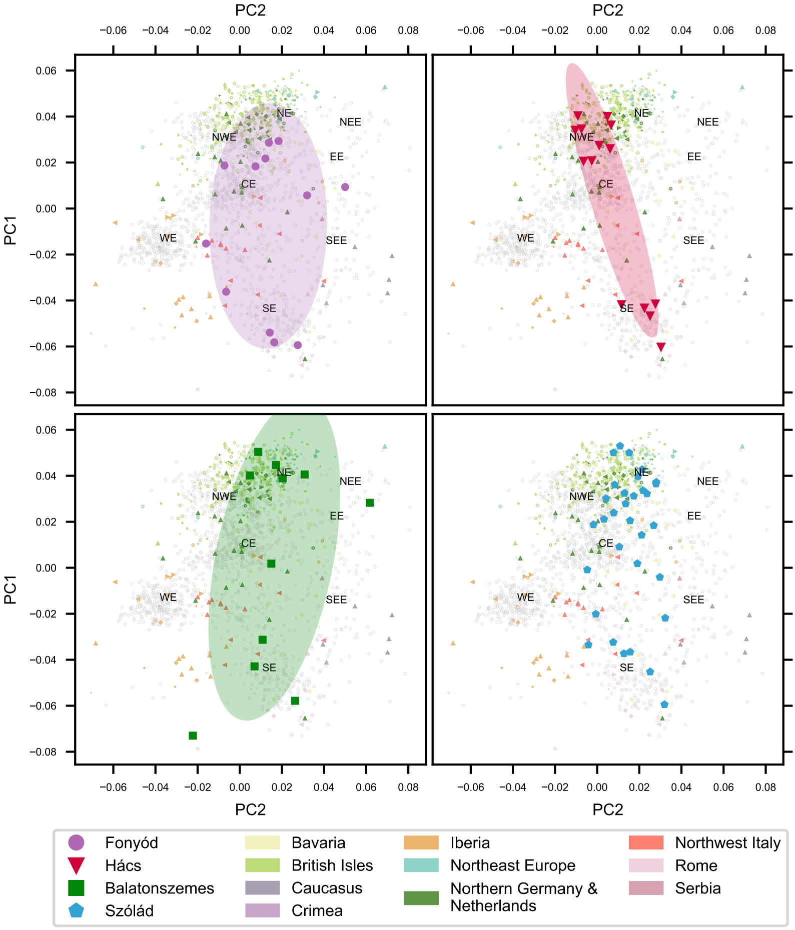
PC2
PC1
NE
NEE
NWE
EE
CE
WE
SEE
SE
Fonyód
Hács
Balatonszemes
Szólád
Bavaria
British Isles
Caucasus
Crimea
Iberia
Northeast Europe
Northern Germany & Netherlands
Northwest Italy
Rome
Serbia

**Figure S1:** Procrustes PCA of 561 individuals[36,41–45,52,69,70] transformed on to a PCA with 1,385 modern European individuals from the POPRES dataset[35] using pseudohaploid genotype calls from 328,670 SNPs with individuals. Penecontemporaneous reference individuals are colored in pastel colors based on their region, while modern individuals from POPRES are colored in grey. Each of the four Lake Balaton communities are plotted in one quadrant with covariance confidence ellipses with radii corresponding to 1.5 standard deviations overlaying the PCA. **a.** Fonyód, **b.** Hács, **c.** Balatonszemes, and **d.** Szólád.

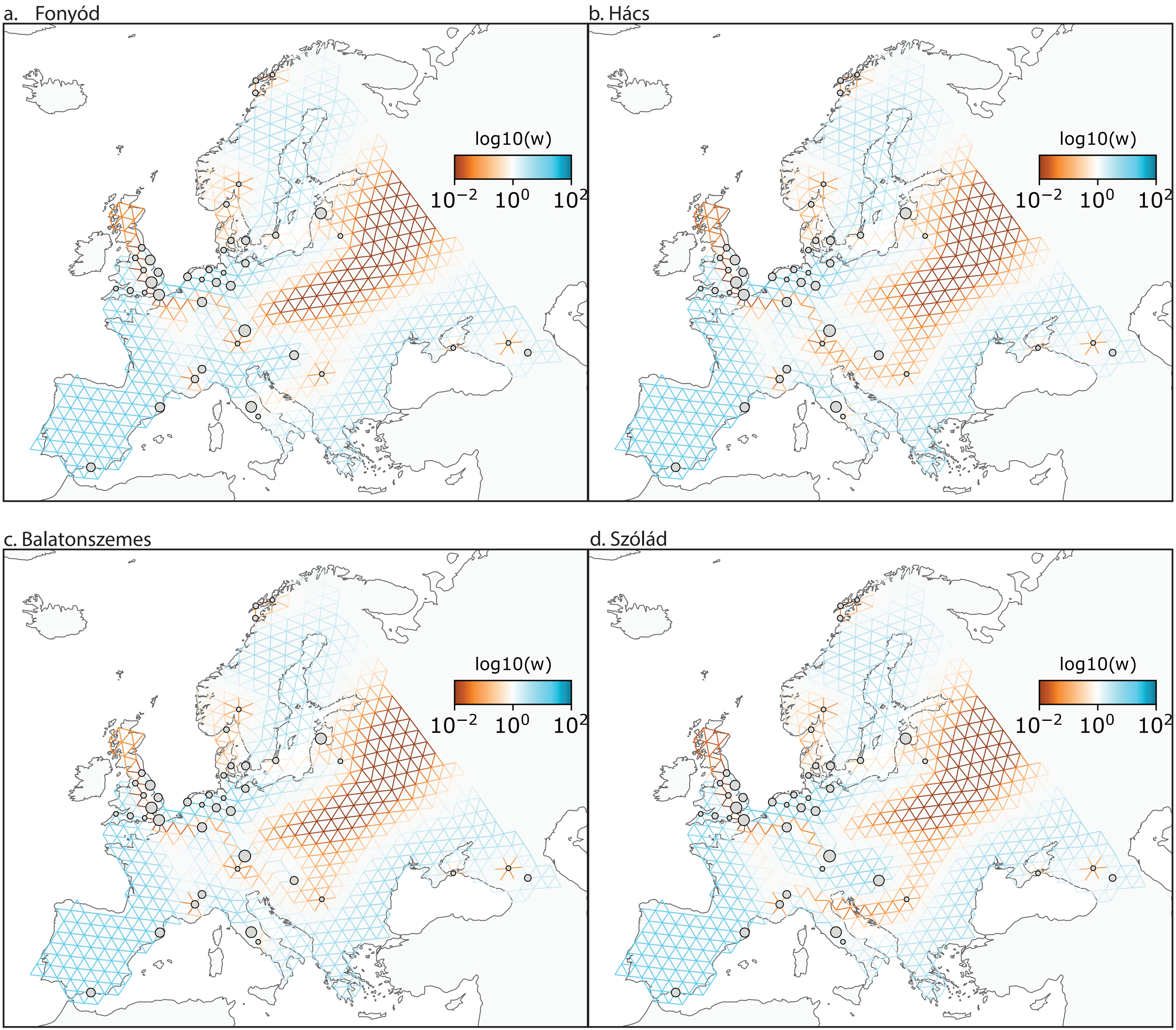
a. Fonyód
b. Hács
c. Balatonszemes
d. Szólád
log10(w)
$10^{-2}$
$10^{0}$
$10^{2}$

**Figure S2:** FEEMS[70] plots estimating gene flow between one Lake Balaton community and the other analyzed 4th-8th century communities. In each quadrant, a different community is analyzed. **a.** Fonyód, **b.** Hács, **c.** Balatonszemes, and **d.** Szólád.

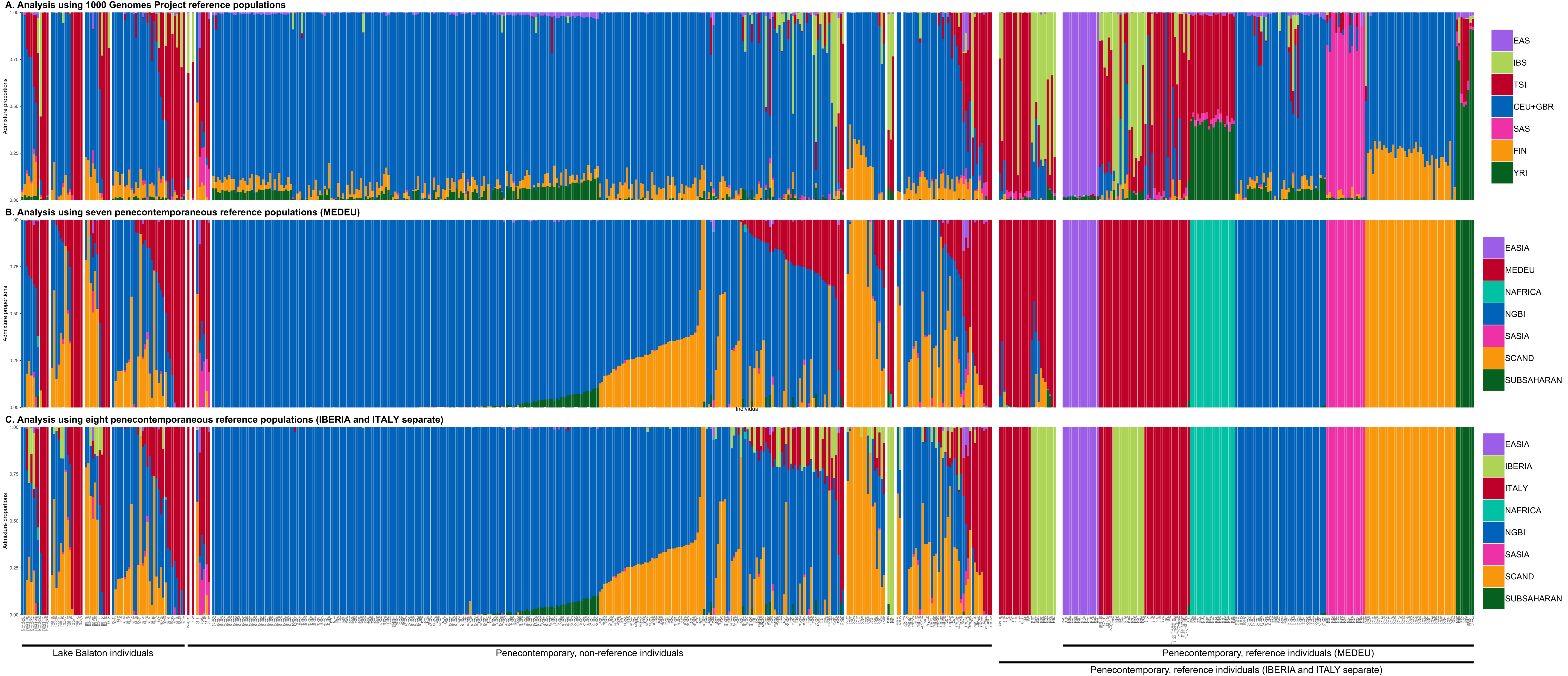
A. Analysis using 1000 Genomes Project reference populations
EAS
IBS
TSI
CEU+GBR
SAS
FIN
YRI
B. Analysis using seven penecontemporaneous reference populations (MEDEU)
EASIA
MEDEU
NAFRICA
NGBI
SASIA
SCAND
SUBSAHARAN
C. Analysis using eight penecontemporaneous reference populations (IBERIA and ITALY separate)
EASIA
IBERIA
ITALY
NAFRICA
NGBI
SASIA
SCAND
SUBSAHARAN
Admixture proportions
Individual
Lake Balaton individuals
Penecontemporary, non-reference individuals
Penecontemporary, reference individuals (MEDEU)
Penecontemporary, reference individuals (IBERIA and ITALY separate)

**Figure S3:** Supervised ancestry proportions from 622 ancient individuals. **a.** using 1000 Genomes populations as references[39] with CEU+GBR referring to a merger of “Northern Europeans from Utah” (CEU) and “British in England and Scotland” (GBR), FIN refers to “Finnish in Finland”, IBS refers to “Iberian populations in Spain”, and TSI refers to “Tuscans from Italy”; EAS refers to the East Asian super-population, SAS refers to the South Asian super-population, and YRI refers to “Yoruba in Ibadan, Nigeria”. **b.** penecontemporaneous individuals to form reference populations[41–51] with MEDEU referring to individuals from Italy and Iberia (Mediterranean Europe), NGBI refers to individuals from what are now northern Germany and Britain, SCAND refers to individuals from what are now Scandinavia/Estonia, EASIA refers to individuals from what is now Hanben, Taiwan, NAFRICA refers to individuals from what is now Sudan, SASIA refers to individuals from Roopkund Lake in what is now India, and SUBSAHARAN refers to individuals from sub-Saharan Africa. Individuals with <0.1× coverage were excluded from this analysis. Individuals are sorted based on increasing MEDEU and then by decreasing NGBI. **c.** penecontemporaneous individuals but with the MEDEU panel replaced by separate IBERIA and ITALY panels. Lake Balaton individuals are presented on the right and are sorted by site, by increasing MEDEU, and then by decreasing NGBI. Penecontemporary individuals are divided by whether or not they are used as reference individuals. Within the non-reference category, individuals are sorted by original publication, by increasing MEDEU, and then by decreasing NGBI. The 25 individuals that are non-reference in **b.** and reference in **c**. are sorted by original publication, by increasing MEDEU, and then by decreasing NGBI. Lastly, the reference individuals (in both **b.** and **c.**) are sorted based on the panel they are included in.

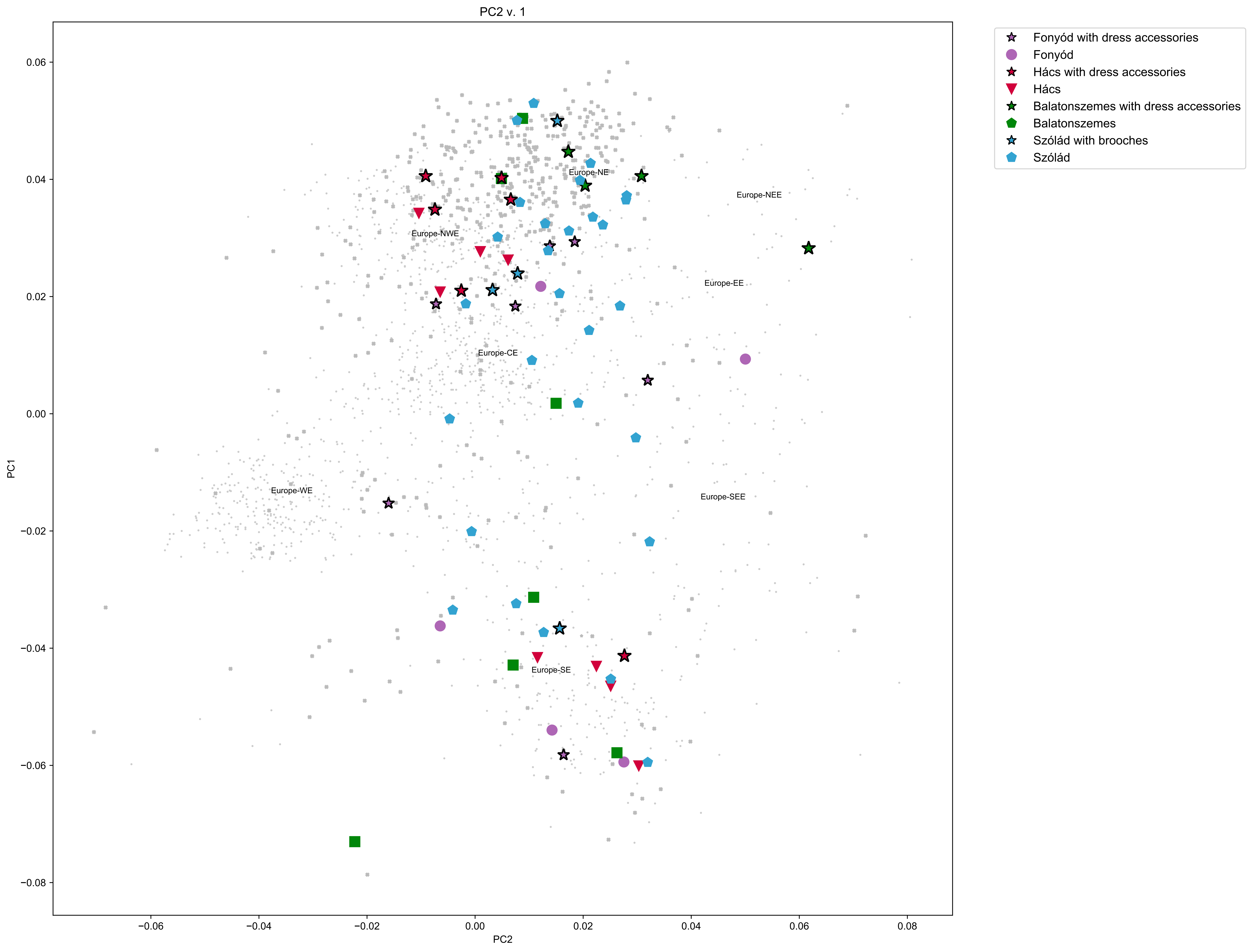
PC2 v. 1
Fonyód with dress accessories
Fonyód
Hács with dress accessories
Hács
Balatonszemes with dress accessories
Balatonszemes
Szólád with brooches
Szólád
Europe-NE
Europe-NEE
Europe-NWE
Europe-EE
Europe-CE
Europe-WE
Europe-SEE
Europe-SE
PC1
PC2
0.06
0.04
0.02
0.00
−0.02
−0.04
−0.06
−0.08
0.08

**Figure S4:** Procrustes PCA of 561 individuals[36,41–45,52,69,70] transformed on to a PCA with 1,385 modern European individuals from the POPRES dataset[35] using pseudohaploid genotype calls from 328,670 SNPs with individuals. Individuals from the Lake Balaton communities are plotted in color, while penecontemporaneous reference individuals and modern individuals from POPRES are both colored in grey. Individuals from Fonyód, Hács, and Balatonszemes who were buried with dress accessories and individuals from Szólád who were buried with brooches are plotted here as stars.

**Supplemental Table S4.** Results from logistic regression analyses

| Sampleset | Sample Size | Variable | Coefficients | | | Null Deviance | Residual Deviance | Overall P value | Hosmer and Lemeshow's $R^2$ | Power ($\alpha$ = 0.05) | If significant, which PC is greater? |
|---|---|---|---|---|---|---|---|---|---|---|---|
| | | | Intercept | PC1 | PC2 | | | | | | |
| Genetic females from Fonyód | 8 | Dress acc. | -52.1679 | 59.62747 | 74.8872 | 8.997362 | 4.047015 | 0.084148 | 0.5502 | NaN | |
| Genetic females from Hács | 8 | Dress acc. | -2201.48 | 1188.605 | 3046.947 | 8.997362 | 8.94E-10 | 0.011124 | 1 | NaN | PC2 |
| Genetic females from Bal. | 7 | Dress acc. | -54.7588 | 75.14036 | 24.03433 | 9.560713 | 4.12E-10 | 0.008393 | 1 | NaN | PC1 |
| Genetic female adults from F/H/B | 20 | Dress acc. | -5.90111 | 6.464049 | 6.728813 | 25.89787 | 16.00929 | 0.007124 | 0.38183 | 0.163564 | PC2 |
| Genetic females from F/H/B | 23 | Dress acc. | -6.47656 | 6.785111 | 7.650034 | 28.26715 | 16.1942 | 0.00239 | 0.427102 | 0.131037 | PC2 |
| Genetic females from Szólád | 11 | Brooches | -0.64459 | 2.342425 | -2.38805 | 14.4206 | 13.30969 | 0.573813 | 0.077036 | 0.050147 | |
| Individuals from Fonyód with preserved skulls | 10 | ACD | 27.69334 | 193.2335 | -258.944 | 13.46023 | 5.38E-10 | 0.001194 | 1 | 0.05 | PC1 |

#### SI References


1. Bierbrauer, V. (2004). Die Keszthely-Kultur und die romanische Kontinuität in Westungarn (5.--8. Jh.). Neue Überlegungen zu einem alten Problem. In Von Sachsen bis Jerusalem. Menschen und Institutionen im Wandel der Zeit. Festschrift für Wolfgang Giese zum 65. Geburtstag, H. Seibert and G. Thoma, eds. (Herbert Utz Verlag), pp. 51–72.

2. Heinrich-Tamáska, O. (2007). Bemerkungen zur Transformation spätantiker Strukturen in Pannonien am Beispiel von Keszthely-Fenékpuszta. Acta Archaeologica Carpathica *42*, 199–229.

3. Schilling, L. (2011). Bestattungen und Gräberfelder von der Spätantike bis zum Frühmittelalter in und um die spätrömische Befestigung von Tác/Gorsium (4.--8. Jh.). In Keszthely-Fenékpuszta im Kontext spätantiker Kontinuitätforschung zwischen Noricum und Moesia. Castellum Pannonicum Pelsonense 2, O. Heinrich-Tamáska, ed. (Leidorf), pp. 381–396.

4. Kiss, A. (1995). Das germanische Gräberfeld von Hács-Béndekpuszta (Westungarn) aus dem 5.--6. Jahrhundert. Acta Antiquae Academiae Scientiarum Hungaricae *36*, 275–342.

5. Bollók, Á. (2016). A fifth-century scriptural amulet from Hács-Béndekpuszta in its Mediterranean context. In Between Byzantium and the Steppe: Archaeological and Historical Studies in Honour of Csanád Bálint on the Occasion of His 70th Birthday, Á. Bollók, G. Csiky, and T. Vida, eds. (Institute of Archaeology, Research Centre for the Humanities, Hungarian Academy of Sciences), pp. 31–61.

6. Bondár, M., Honti, S., Márkus, G., and Németh, P.G. (2007). Balatonszemes-Szemesi berek. In Gördülő idő: régészeti feltárások az M7-es autópálya Somogy megyei szakaszán Zamárdi és Ordacsehi között, K. Belényesy, S. Honti, and V. Kiss, eds. (Somogy Megyei Múzeumok Igazgatósága), pp. 123–135.

7. Straub, P. (2008). Adalékok a Balaton környéki 5. századi temetők Felső-Duna vidéki kapcsolatához. Zalai Múzeum *17*, 189–207.

8. Miháczi-Pálfi, A. (2017). Form-und herstellungstechnische Analyse der Bügelfibeln von Balatonszemes aus dem dritten Viertel des 5. Jahrhunderts. ANTÆUS: Communicationes Ex Instituto Archaeologico Academiae Scientiarum Hungaricae *35-36*, 67–89.

9. Pejrani Baricco, L., and Ratto, S. (2014). L'inattesa scoperta di una chiesa paleocristiana. Rivista MuseoTorino *7*, 10–13.

10. Giostra, C., Bedini, E., Caramelli, D., Mallegni, F., and Pejrani Baricco, L. (2012). Per una conoscenza dei Longobardi in Italia: primi risultati delle analisi genetiche su individui provenienti da necropoli del Piemonte. In VI Congresso Nazionale di Archeologia Medievale, L'Aquila, 12-15 settembre 2012 (ITA), pp. 448–453.

11. Pejrani Baricco, L. (2017). Bardonecchia (Torino), necropoli di ambito merovingio. In Longobardi. Un popolo che cambia la storia, catalogo della mostra (Pavia – Napoli – San Pietroburgo, 2017-2018), G. P. Brogiolo, F. Marazzi, and C. Giostra, eds. (Catalogo di mostra internazionale allestita presso il Castello Visconteo di Pavia, il Museo Archeologico Nazionale di Napoli e l'Ermitage di San Pietroburgo.), pp. 76–77.

12. Gilbert, M.T.P., Bandelt, H.-J., Hofreiter, M., and Barnes, I. (2005). Assessing ancient DNA studies. Trends Ecol. Evol. *20*, 541–544.

13. Willerslev, E., and Cooper, A. (2005). Ancient DNA. Proc. Biol. Sci. *272*, 3–16.

14. Pinhasi, R., Fernandes, D., Sirak, K., Novak, M., Connell, S., Alpaslan-Roodenberg, S., Gerritsen, F., Moiseyev, V., Gromov, A., Raczky, P., et al. (2015). Optimal Ancient DNA Yields from the Inner Ear Part of the Human Petrous Bone. PLoS One *10*, e0129102.

15. Dabney, J., Knapp, M., Glocke, I., Gansauge, M.-T., Weihmann, A., Nickel, B., Valdiosera, C., García, N., Pääbo, S., Arsuaga, J.-L., et al. (2013). Complete mitochondrial genome sequence of a Middle Pleistocene cave bear reconstructed from ultrashort DNA fragments. Proc Natl Acad Sci USA *110*, 15758–15763.

16. Meyer, M., and Kircher, M. (2010). Illumina sequencing library preparation for highly multiplexed target capture and sequencing. Cold Spring Harb. Protoc. *2010*, db.prot5448.

17. Kircher, M., Sawyer, S., and Meyer, M. (2012). Double indexing overcomes inaccuracies in multiplex sequencing on the Illumina platform. Nucleic Acids Res. *40*, e3.

18. Fu, Q., Meyer, M., Gao, X., Stenzel, U., Burbano, H.A., Kelso, J., and Pääbo, S. (2013). DNA analysis of an early modern human from Tianyuan Cave, China. Proc. Natl. Acad. Sci. U. S. A. *110*, 2223–2227.

19. Haak, W., Lazaridis, I., Patterson, N., Rohland, N., Mallick, S., Llamas, B., Brandt, G., Nordenfelt, S., Harney, E., Stewardson, K., et al. (2015). Massive migration from the steppe was a source for Indo-European languages in Europe. Nature *522*, 207–211.

20. Mathieson, I., Lazaridis, I., Rohland, N., Mallick, S., Patterson, N., Roodenberg, S.A., Harney, E., Stewardson, K., Fernandes, D., Novak, M., et al. (2015). Genome-wide patterns of selection in 230 ancient Eurasians. Nature *528*, 499–503.

21. Gansauge, M.-T., and Meyer, M. (2013). Single-stranded DNA library preparation for the sequencing of ancient or damaged DNA. Nat. Protoc. *8*, 737–748.

22. Gansauge, M.-T., and Meyer, M. (2019). A Method for Single-Stranded Ancient DNA Library Preparation. Methods Mol. Biol. *1963*, 75–83.

23. Rohland, N., Harney, E., Mallick, S., Nordenfelt, S., and Reich, D. (2015). Partial uracil-DNA-glycosylase treatment for screening of ancient DNA. Philos. Trans. R. Soc. Lond. B Biol. Sci. *370*, 20130624.

24. Rohland, N., Glocke, I., Aximu-Petri, A., and Meyer, M. (2018). Extraction of highly degraded DNA from ancient bones, teeth and sediments for high-throughput sequencing. Nat. Protoc. *13*, 2447–2461.

25. Gansauge, M.-T., Gerber, T., Glocke, I., Korlevic, P., Lippik, L., Nagel, S., Riehl, L.M., Schmidt, A., and Meyer, M. (2017). Single-stranded DNA library preparation from highly degraded DNA using T4 DNA ligase. Nucleic Acids Res. *45*, e79.

26. Gansauge, M.-T., Aximu-Petri, A., Nagel, S., and Meyer, M. (2020). Manual and automated preparation of single-stranded DNA libraries for the sequencing of DNA from ancient biological remains and other sources of highly degraded DNA. Nat. Protoc. *15*, 2279–2300.

27. DeAngelis, M.M., Wang, D.G., and Hawkins, T.L. (1995). Solid-phase reversible immobilization for the isolation of PCR products. Nucleic Acids Res. *23*, 4742–4743.

28. Kircher, M. (2012). Analysis of high-throughput ancient DNA sequencing data. Methods Mol. Biol. *840*, 197–228.

29. Danecek, P., Bonfield, J.K., Liddle, J., Marshall, J., Ohan, V., Pollard, M.O., Whitwham, A., Keane, T., McCarthy, S.A., Davies, R.M., et al. (2021). Twelve years of SAMtools and BCFtools. Gigascience *10*. 10.1093/gigascience/giab008.

30. McKenna, A., Hanna, M., Banks, E., Sivachenko, A., Cibulskis, K., Kernytsky, A., Garimella, K., Altshuler, D., Gabriel, S., Daly, M., et al. (2010). The Genome Analysis Toolkit: a MapReduce framework for analyzing next-generation DNA sequencing data. Genome Res. *20*, 1297–1303.

31. Ginolhac, A., Rasmussen, M., Gilbert, M.T.P., Willerslev, E., and Orlando, L. (2011). mapDamage: testing for damage patterns in ancient DNA sequences. Bioinformatics *27*, 2153–2155.

32. Korneliussen, T.S., Albrechtsen, A., and Nielsen, R. (2014). ANGSD: Analysis of Next Generation Sequencing Data. BMC Bioinformatics *15*, 356.

33. Renaud, G., Slon, V., Duggan, A.T., and Kelso, J. (2015). Schmutzi: estimation of contamination and endogenous mitochondrial consensus calling for ancient DNA. Genome Biol. *16*, 224.

34. Gnecchi-Ruscone, G.A., Szécsényi-Nagy, A., Koncz, I., Csiky, G., Rácz, Z., Rohrlach, A.B., Brandt, G., Rohland, N., Csáky, V., Cheronet, O., et al. (2022). Ancient genomes reveal origin and rapid trans-Eurasian migration of 7th century Avar elites. Cell *185*, 1402–1413.e21.

35. Nelson, M.R., Bryc, K., King, K.S., Indap, A., Boyko, A.R., Novembre, J., Briley, L.P., Maruyama, Y., Waterworth, D.M., Waeber, G., et al. (2008). The Population Reference Sample, POPRES: a resource for population, disease, and pharmacological genetics research. Am. J. Hum. Genet. *83*, 347–358.

36. Veeramah, K.R., Rott, A., Groß, M., van Dorp, L., López, S., Kirsanow, K., Sell, C., Blöcher, J., Wegmann, D., Link, V., et al. (2018). Population genomic analysis of elongated skulls reveals extensive female-biased immigration in Early Medieval Bavaria. Proc. Natl. Acad. Sci. U. S. A. *115*, 3494–3499.

37. Price, A.L., Patterson, N.J., Plenge, R.M., Weinblatt, M.E., Shadick, N.A., and Reich, D. (2006). Principal components analysis corrects for stratification in genome-wide association studies. Nat. Genet. *38*, 904–909.

38. Patterson, N., Price, A.L., and Reich, D. (2006). Population structure and eigenanalysis. PLoS Genet. *2*, e190.

39. 1000 Genomes Project Consortium, Auton, A., Brooks, L.D., Durbin, R.M., Garrison, E.P., Kang, H.M., Korbel, J.O., Marchini, J.L., McCarthy, S., McVean, G.A., et al. (2015). A global reference for human genetic variation. Nature *526*, 68–74.

40. Rubinacci, S., Ribeiro, D.M., Hofmeister, R.J., and Delaneau, O. (2021). Efficient phasing and imputation of low-coverage sequencing data using large reference panels. Nat. Genet. *53*, 120–126.

41. Margaryan, A., Lawson, D.J., Sikora, M., Racimo, F., Rasmussen, S., Moltke, I., Cassidy, L.M., Jørsboe, E., Ingason, A., Pedersen, M.W., et al. (2020). Population genomics of the Viking world. Nature *585*, 390–396.

42. Gretzinger, J., Sayer, D., Justeau, P., Altena, E., Pala, M., Dulias, K., Edwards, C.J., Jodoin, S., Lacher, L., Sabin, S., et al. (2022). The Anglo-Saxon migration and the formation of the early English gene pool. Nature *610*, 112–119.

43. Schiffels, S., Haak, W., Paajanen, P., Llamas, B., Popescu, E., Loe, L., Clarke, R., Lyons, A., Mortimer, R., Sayer, D., et al. (2016). Iron Age and Anglo-Saxon genomes from East England reveal British migration history. Nat. Commun. *7*, 1–9.

44. Antonio, M.L., Gao, Z., Moots, H.M., Lucci, M., Candilio, F., Sawyer, S., Oberreiter, V., Calderon, D., Devitofranceschi, K., Aikens, R.C., et al. (2019). Ancient Rome: A genetic crossroads of Europe and the Mediterranean. Science *366*, 708–714.

45. Olalde, I., Mallick, S., Patterson, N., Rohland, N., Villalba-Mouco, V., Silva, M., Dulias, K., Edwards, C.J., Gandini, F., Pala, M., et al. (2019). The genomic history of the Iberian Peninsula over the past 8000 years. Science *363*, 1230–1234.

46. Harney, É., Nayak, A., Patterson, N., Joglekar, P., Mushrif-Tripathy, V., Mallick, S., Rohland, N., Sedig, J., Adamski, N., Bernardos, R., et al. (2019). Ancient DNA from the skeletons of Roopkund Lake reveals Mediterranean migrants in India. Nat. Commun. *10*, 1–10.

47. Sirak, K.A., Fernandes, D.M., Lipson, M., Mallick, S., Mah, M., Olalde, I., Ringbauer, H., Rohland, N., Hadden, C.S., Harney, É., et al. (2021). Social stratification without genetic differentiation at the site of Kulubnarti in Christian Period Nubia. Nat. Commun. *12*, 1–14.

48. Skoglund, P., Thompson, J.C., Prendergast, M.E., Mittnik, A., Sirak, K., Hajdinjak, M., Salie, T., Rohland, N., Mallick, S., Peltzer, A., et al. (2017). Reconstructing Prehistoric African Population Structure. Cell *171*, 59–71.e21.

49. Wang, K., Goldstein, S., Bleasdale, M., Clist, B., Bostoen, K., Bakwa-Lufu, P., Buck, L.T., Crowther, A., Dème, A., McIntosh, R.J., et al. (2020). Ancient genomes reveal complex patterns of population movement, interaction, and replacement in sub-Saharan Africa. Science Advances *6*, eaaz0183.

50. Wang, C.-C., Yeh, H.-Y., Popov, A.N., Zhang, H.-Q., Matsumura, H., Sirak, K., Cheronet, O., Kovalev, A., Rohland, N., Kim, A.M., et al. (2021). Genomic insights into the formation of human populations in East Asia. Nature *591*, 413–419.

51. Prendergast, M.E., Lipson, M., Sawchuk, E.A., Olalde, I., Ogola, C.A., Rohland, N., Sirak, K.A., Adamski, N., Bernardos, R., Broomandkhoshbacht, N., et al. (2019). Ancient DNA reveals a multistep spread of the first herders into sub-Saharan Africa. Science *365*, eaaw6275.

52. Amorim, C.E.G., Vai, S., Posth, C., Modi, A., Koncz, I., Hakenbeck, S., La Rocca, M.C., Mende, B., Bobo, D., Pohl, W., et al. (2018). Understanding 6th-century barbarian social organization and migration through paleogenomics. Nat. Commun. *9*, 3547.

53. Ensor, B.E. (2013). The Archaeology of Kinship: Advancing Interpretation and Contributions to Theory (University of Arizona Press).

54. Hummer, H. (2018). Visions of Kinship in Medieval Europe (Oxford University Press).

55. Brück, J. (2021). Ancient DNA, kinship and relational identities in Bronze Age Britain. Antiquity *95*, 228–237.

56. Rácz, Z. (2016). Zwischen Hunnen- und Gepidenzeit. Frauengräber aus dem 5. Jahrhundert im Karpatenbecken. Acta Archaeologica Academiae Scientiarum Hungaricae *67*, 301–359.

57. Rácz, Z. (2020). Who Were the Gepids and Ostrogoths on the Middle Danube in the 5th Century? An Archaeological Perspective. In The Migration Period between the Oder and the Vistula (2 vols) (Brill), pp. 771–789.

58. Hakenbeck, S.E., Evans, J., Chapman, H., and Fóthi, E. (2017). Practising pastoralism in an agricultural environment: An isotopic analysis of the impact of the Hunnic incursions on Pannonian populations. PLoS One *12*, e0173079.

59. Katoh, K., Misawa, K., Kuma, K., and Miyata, T. (2002). MAFFT: a novel method for rapid multiple sequence alignment based on fast Fourier transform. Nucleic Acids Res. *30*, 3059–3066.

60. Katoh, K., and Standley, D.M. (2013). MAFFT Multiple Sequence Alignment Software Version 7: Improvements in Performance and Usability. Mol. Biol. Evol. *30*, 772–780.

61. van Oven, M., and Kayser, M. (2009). Updated comprehensive phylogenetic tree of global human mitochondrial DNA variation. Hum. Mutat. *30*, E386–E394.

62. Weir, B.S., and Cockerham, C.C. (1984). Estimating F-Statistics for the Analysis of Population Structure. Evolution *38*, 1358–1370.

63. Excoffier, L., Smouse, P.E., and Quattro, J.M. (1992). Analysis of molecular variance inferred from metric distances among DNA haplotypes: application to human mitochondrial DNA restriction data. Genetics *131*, 479–491.

64. Weir, B.S. (1996). Genetic Data Analysis II: Methods for Discrete Population Genetic Data (Sinauer Associates).

65. Raymond, M., and Rousset, F. (1995). An exact test for population differentiation. Evolution *49*, 1280–1283.

66. Goudet, J., Raymond, M., de Meeüs, T., and Rousset, F. (1996). Testing differentiation in diploid populations. Genetics *144*, 1933–1940.

67. Nei, M. (1987). Molecular evolutionary genetics (Columbia University Press).

68. Excoffier, L., and Lischer, H.E.L. (2010). Arlequin suite ver 3.5: a new series of programs to perform population genetics analyses under Linux and Windows. Mol. Ecol. Resour. *10*, 564–567.

69. Damgaard, P. de B., Marchi, N., Rasmussen, S., Peyrot, M., Renaud, G., Korneliussen, T., Moreno-Mayar, J.V., Pedersen, M.W., Goldberg, A., Usmanova, E., et al. (2018). 137 ancient human genomes from across the Eurasian steppes. Nature *557*, 369–374.

70. Marcus, J., Ha, W., Barber, R.F., and Novembre, J. (2021). Fast and flexible estimation of effective migration surfaces. Elife *10*, e61927.
